# The hexokinase “HKDC1” interaction with the mitochondria is essential for hepatocellular carcinoma progression

**DOI:** 10.1101/2021.11.10.468146

**Authors:** Md. Wasim Khan, Alexander R. Terry, Medha Priyadarshini, Grace Guzman, Jose Cordoba-Chacon, Issam Ben-Sahra, Barton Wicksteed, Brian T. Layden

## Abstract

Hepatocellular carcinoma (HCC) is a leading cause of death from cancer malignancies. Recently, hexokinase domain containing 1 (HKDC1), was shown to have significant overexpression in HCC compared to healthy tissue. Using *in vitro* and *in vivo* tools, we examined the role of HKDC1 in HCC progression. Importantly, HKDC1 ablation stops HCC progression by promoting metabolic reprogramming by shifting glucose flux away from the TCA cycle. Next, HKDC1 ablation leads to mitochondrial dysfunction resulting in less cellular energy which cannot be compensated by enhanced glucose uptake. And finally, we show that the interaction of HKDC1 with the mitochondria is essential for its role in HCC progression, and without this mitochondrial interaction mitochondrial dysfunction occurs. In sum, HKDC1 is highly expressed in HCC cells compared to normal hepatocytes, therefore targeting HKDC1, specifically its interaction with the mitochondria, reveals a highly selective approach to target cancer cells in HCC.

## Introduction

Globally, there has been a decline in number of deaths due to cancer malignancies; however, the incidence and mortality due to liver cancer continues to rise (Ryerson et al., 2016). The most common form of liver cancer is hepatocellular carcinoma (HCC), and it is the fifth leading cause of death due to cancer worldwide (Zhou et al., 2021, Orci et al., 2021, Nagaoki et al., 2021). HCC commonly occurs in the settings of chronic liver diseases secondary to a) viral (chronic hepatitis B and C), b) toxic (alcohol and aflatoxin), c) immune (autoimmune hepatitis and primary biliary) and d) metabolic (diabetes and nonalcoholic liver disease) origins (Orci et al., 2021, Nagaoki et al., 2021). The high mortality rates of HCC are ascribed to its aggressiveness and heterogeneity and possibly the accompanying severe liver dysfunction.

Until now, a cure is feasible only if the disease is localized so that complete surgical resection can occur. As the risk of recurrence is very high, there is a need for discoveries of novel targets that can be used for prognostic and diagnostic opportunities in HCC (Bruix and Sherman, 2005). One such possible target is the phenomenon of “metabolic reprogramming” that occurs during carcinogenesis particularly due to the following reasons: 1) metabolism is reprogrammed in cancer, 2) oncogenes drive metabolic reprogramming, 3) gene/protein expression is regulated by altered cancer metabolites and 4) dysregulated metabolic protein/enzymes represent diagnostic and/or prognostic biomarkers (DeBerardinis and Chandel, 2016, Wolpaw and Dang, 2018, Jin et al., 2020, Bao and Wong, 2021). Most cancer cells/types have an over-dependency on glucose, and this dysregulated glucose metabolism can be therapeutically used to target specific vulnerabilities (DeBerardinis and Chandel, 2016, Wolpaw and Dang, 2018). Therefore, targeting pathways that metabolize glucose specifically in HCC may constitute novel approaches in HCC treatment.

Phosphorylation is the first step of glucose catabolism, where hexokinases (HKs) catalyze this critical first step in the regulation of energy metabolism that particularly contributes to the rate of cell growth and proliferation (Middleton, 1990, Wilson, 1995). HK1 is widely and constitutively expressed, whereas HK2 is expressed during embryogenesis and is not widely expressed in adult tissues as HK1 (Middleton, 1990, Wilson, 1995). However, HK2 is selectively overexpressed in different form of cancer types including HCC where it has been shown to act along with partner proteins to induce proliferation and inhibit cell death (Lis et al., 2016, Jin et al., 2017, DeWaal et al., 2018).

Along with a role of HK2 in HCC, a major driver in the development of HCC is hepatocyte mitochondrial dysfunction (via its role in metabolism and cellular survival) (Robey and Hay, 2006, McDonald et al., 2019). There are various possible mechanisms by which altering mitochondrial function augments HCC progression such as i) generation of ROS which enhances carcinogenesis, ii) changes in activities of mitochondrial enzymes involved in TCA cycle (SDH, FH and IDH) leads to accumulation of oncometabolites that aid in oncogenesis and, iii) defects in the mitochondria may lead to faulty apoptotic pathways that regulate cell death (Luo et al., 2020, Srinivasan et al., 2017). HK1 and HK2 have been shown to interact with the mitochondria physically and functionally and play roles in cancer cell survival, in particular for HK2 (Majewski et al., 2004, Pastorino et al., 2002). Thus, HKs are potentially important contributors to HCC progression through their effects at the mitochondria (Robey and Hay, 2006, Smith, 2000). We recently discovered a novel 5th HK, hexokinase domain containing 1 (HKDC1) which has been shown to be overexpressed in certain cancers compared to healthy tissue, with the most significant changes in HCC as compared to other HKs (Chen et al., 2019, Guo et al., 2015, Hayes et al., 2013, Irwin and Tan, 2008, Li and Huang, 2014, Xu et al., 2021, Zhang et al., 2016, Cerami et al., 2012). Collectively, our data indicate that HKDC1 plays a pivotal role in HCC progression, via its action at the mitochondria and therefore can be potentially promising target for therapeutic intervention.

## Materials and Methods

### Animal studies

All animal experiments were approved by the University of Illinois at Chicago Institutional Animal Care and Use Committee, as required by United States Animal Welfare Act, and the NIH’s policy.

For Fig 1D, mice were either fed low fat, cholesterol, and fructose (LFCF, Control, Cat # D09100304) or the high fat, cholesterol, and fructose (HFCF, NASH deit, Cat # D16010101, Research diets, Inc) diets for 24 weeks.

**Fig 1.**
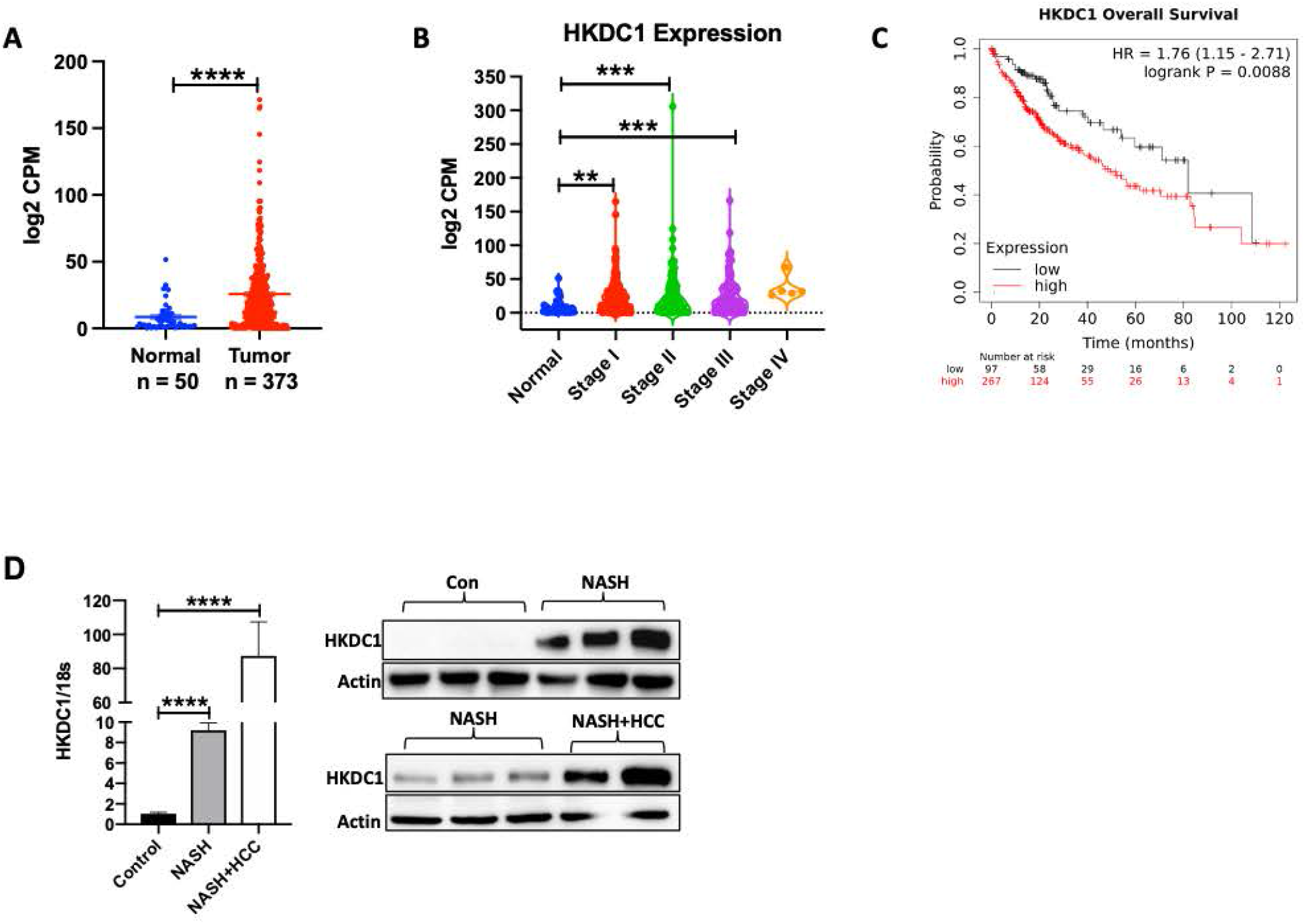
HKDC1 is upregulated in HCC. mRNA expression data of HKDC1 in liver cancer (LIHC) in human patients mined from the TCGA data set is shown for **A)** normal and HCC patient samples where T denotes tumor (n=369) and N=denotes normal surrounding tissue (n=50) and **B)** to show HKDC1 expression in samples from different tumor stages stage I (n=173); stage II (n=87); stage III (n=85); stage IV (n=5); compared to normal tissue (n=50). **C)** Kaplan-Meier survival curves were plotted from TCGA dataset using the website (https://kmplot.com/analysis/) with no restrictions applied; HR=hazard ratio. **D)** HKDC1 mRNA (left panel) and protein expression (right panel) from liver (non-tumor regions) of mice fed control diet (Con) or NASH diet (NASH) over 20 weeks (n=5 for control and NASH; n=3 for NASH+HCC), mice that developed HCC along with NASH are denoted NASH+HCC. Values are mean ± SEM; *p < 0.05; **p < 0.01; ***p < 0.001; ****p < 0.0001 by student’s t-test (for 1A) or one-way ANOVA (for 1B, D).

#### DEN model

For diethylnitrosamine (DEN) induced hepatocarcinogenesis, HKDC1 floxed mice were crossed with Albumin-Cre mice (Jackson Lab, Stock No: 003574) (Postic et al., 1999) to create a liver specific HKDC1 knockout mouse model (HKDC1-LKO). DEN-induced hepatocarcinogenesis was performed by intraperitoneal injection of (25 mg/kg body weight) in 14 days old male mice (Tolba et al., 2015). Livers from 10 month-old treated mice were analyzed macroscopically for tumor lesions. Livers were then collected fixed in 10% formalin for 18 h and subsequently preserved with 70% ethanol. Fixed tissues were then processed and embedded in paraffin. Paraffin embedded tissues were processed and 5 μM slides were prepared for BrdU and Ki67 staining.

#### Xenograft model

HepG2 cells were deleted for HKDC1 by CRISPR/Cas9 (described below). Male athymic nude mice (5 weeks old) were subcutaneously injected into the right flank with 1×10^6^ (in 100 μL) HepG2 cells-expressing either empty vector (EV) or HKDC1-KO cells. When tumors were palpable (∼100 mm^3^), tumor size was monitored twice per week and measurements were taken with a Vernier caliper till 16 weeks post implantation. Tumor volume was calculated by the formula (a×b^2^)/2 where a and b are length and breadth, respectively. End points were reached, and mice were sacrificed once the tumor size measured 2 cm.

For the inducible shRNA mediated HKDC1 knockdown experiments, modulated Hep3B2 cells were grown in culture before collection for injection subcutaneously into the right flanks of each mouse (1×10^6^ in 100 μL per site). Once tumors were ∼65 mm^3^ in volume, mice from each group were given a treatment with a (200 mg/kg) doxycycline diet (Bioserve) for 7 days. Tumor volume was calculated by the formula (a×b^2^)/2 where a and b are length and breadth, respectively. End points were reached, and mice were sacrificed once the tumor size measured 2 cm.

### Cell culture and reagents

HepG2, Huh7, Hep3B2, HCT116, cells were grown in Eagle’s Minimum Essential Medium (EMEM; ATCC; 30-2003) supplemented with 10% fetal bovine serum (FBS; Sigma; F0926) and 1% penicillin-streptomycin (Gibco; 15070063). SNU-475, MCF10, MDA-MB-231, BT-20, T47D, RC45 and RC26B cells were maintained in Roswell Park Memorial Institute Medium (RPMI; ATCC; 30-2001) supplemented with 10% fetal bovine serum (FBS; Sigma; F0926) and 1% penicillin-streptomycin (Gibco; 15070063). All chemicals were from Sigma-Aldrich (St Louis, MO, US) unless stated otherwise.

### Cell lysis and immunoblotting

Cells were rinsed with ice-cold PBS before lysis in buffer containing 50 mM Tris-HCl (pH 7.4), 100 mM NaCl, 1 mM EDTA, 1 mM EGTA and 1% Triton X-100 with protease/phosphatase inhibitor (Roche) mixture. The soluble fractions of cell lysates were isolated by centrifugation at 12000 g for 15 min at 4 °C. Protein concentration was determined using Bradford reagent (Bio-Rad Laboratories, Hercules, CA, USA). 30 μg of denatured proteins were separated by SDS-PAGE (Mini-PROTEAN TGX Gels 10%, Bio-Rad Laboratories) and transferred to 0.45 μm nitrocellulose membranes. Membranes were blocked with 5% nonfat, dried milk in Tris-buffered saline with 0.1% Tween-20 for 1 h at 25 °C, then incubated with primary antibodies overnight at 4 °C washed and incubated with secondary antibodies for 1 h at 25 °C. After washing, Clarity Western ECL Substrate (Bio-Rad) was added, and the light signal was detected and analyzed using a ChemiDoc MP and Image Lab Ver 6.0 (Bio-Rad). List of all antibodies used with dilutions is included in Supplementary Table 2.

### Cell Proliferation assay

Proliferation rates were assessed in cell lines by proliferation curve analysis. Cells were plated on 96-cm clear bottom black well. For proliferation curves, 1 day of growth was allowed before the first cell count, and thereafter, counts were performed every other day for 3-4 total counts. Growth medium was replaced every two days to ensure no nutrient depletion would. Total viable cell numbers of individual wells were determined by staining the cells with Hoechst and propidium iodide (PI) and counting on the Celigo Imaging cytometer (Nexcelom Bioscience LLC, MA, USA). Cells stained with PI were counted as dead and subtracted from total cells to get viable cell number. Cell numbers for each time point were averaged and plotted by scatter plot with standard error of the mean.

### Crispr-Cas9 mediated HKDC1 knockout

HKDC1 sgRNA CRISPR All-in-One Lentivirus (Human) (Cat No. 234181110603) was purchased from ABM and HepG2 cells (70% confluent) were infected at an MOI of 50 (virus titer >1×10⁷IU/ml). Cells were allowed to grow for 2 days post infection after which they were selected with puromycin (1μg/ml) for 7 days. Selected cells were then used to perform a single cell colony formation in a 96 well plate. When colonies were formed (roughly 21 days), the cells from each well were subcultured for 3 generations more. HKDC1 knockout was confirmed in these clones by qPCR and immunoblot and then 3 individual clones were pooled together. Genome edit was confirmed by genomic sequencing in this pool and further HKDC1 knockout was confirmed by qPCR and western blotting.

### Colony formation

200 cells were plated onto 6 well culture plates and allowed to grow for 21 days with media changed every 3 days. Colonies were fixed with glutaraldehyde (6.0% v/v), stained with crystal violet (0.5% w/v) and imaged.

### Glucose consumption

50μl media was collected from cells growing in 6 well plates at 70% confluency at 0, 2, 4, 6 and 16 h and stored at −80C. Glucose was measured in these aliquots with the Biosen R-Line lactate and glucose analyzer (EKF Diagnostics, TX, USA). To calculate glucose consumption, glucose remaining in the media at the indicated time-points was subtracted from the original glucose concentration at 0 h.

### Glucose uptake

Glucose uptake experiments were performed using 2-NBDG (2-(N-(7-nitrobenz-2-oxa-1,3-diazol-4-yl)amino)-2-deoxyglucose) (Invitrogen, Carlsbad, CA, USA) according to the manufacturer’s protocol. Briefly, cells were plated in a 96-well black clear bottom plate (Brand, Wertheim, Germany). After treatment, cells were washed three times with 1× PBS at room temperature and incubated for 30 min in zero glucose DMEM containing 75 μM 2-NBDG. Cells were then washed with ice-cold PBS three times. To each well, 200 μl of PBS was added and the relative fluorescence was measured in a fluorimeter (Synergy H1 multimode microplate reader; Biotek (Winooski, VT, USA); excitation 485 nm, emission 535 nm). The assay was normalized to total cellular protein.

### Hexokinase activity

Hexokinase activity was assayed in cell homogenates using the Hexokinase activity kit (MAK091, Sigma, St. Louis, MO, USA) according to the manufacturer’s instructions. Activity was normalized to the amount of protein in the sample.

### Mitochondrial Complex Assays

Activities for mitochondrial Complex I, SDH and Complex III were performed with commercially available kits (Catalog # K968, K660 and K520; Biovision, CA, US) according to the manufacturer’s protocols.

### Metabolite measurements and analysis

For steady state metabolomics, cultured cells were washed twice with PBS at room temperature and incubated with normal growth media for 2 hours. For isotopic labeling analysis, cells were washed twice with warm PBS and incubated in medium containing 5.5mm of [U-^13^C] glucose, 2mM pyruvate and 2mM glutamine for 4 hours. After incubation, cells were washed twice with ice-cold saline and metabolites were extracted with 1ml of ice-cold 80% methanol per plate. Cells were scraped and samples were frozen in liquid nitrogen and thawed in a 37 °C water bath three times. Following this, cells were centrifuged at 20,000g for 15 min at 4 °C. The supernatant was transferred to fresh tubes and dried. Samples were resuspended in 10 µl per 150,000 cells. Samples were analyzed by high-performance liquid chromatography and high-resolution mass spectrometry, and tandem mass spectrometry (HPLC-MS/MS). Specifically, the system consisted of a Thermo Q-Exactive in line with an electrospray source and an Ultimate3000 (Thermo) series HPLC consisting of a binary pump, degasser and auto-sampler outfitted with an Xbridge amide column (Waters; dimensions 4.6 mm × 100 mm and a 3.5-µm particle size). The mobile phase A contained 95% (v/v) water, 5% (v/v) acetonitrile, 20 mM ammonium hydroxide, 20 mM ammonium acetate, pH 9.0; B was 100% acetonitrile. The gradient was as follows: 0–1 min, 15% A; 18.5 min, 76% A; 18.5–20.4 min, 24% A; 20.4–20.5 min, 15% A; 20.5–28 min, 15% A with a flow rate of 400 μl/min. The capillary of the ESI source was set to 275 °C, with sheath gas at 45 arbitrary units, auxiliary gas at 5 arbitrary units and the spray voltage at 4.0 kV. In positive/negative polarity switching mode, an m/z scan range from 70 to 850 was chosen and MS1 data was collected at a resolution of 70,000. The automatic gain control (AGC) target was set at 1 × 10^6^ and the maximum injection time was 200 ms. The top 5 precursor ions were subsequently fragmented, in a data-dependent manner, using the higher energy collisional dissociation cell set to 30% normalized collision energy in MS2 at a resolution power of 17,500. Sample volumes of 150,000 cells in 10 μl were injected. Data acquisition and analysis were carried out by Xcalibur 4.0 software and Tracefinder 2.1 software, respectively (both from Thermo Fisher Scientific).

### Oxygen consumption rate and extracellular acidification rate

Oxygen consumption rate (OCR) and extracellular acidification rate (ECAR) were measured using an XFe96 extracellular flux analyzer (Seahorse Bioscience). Basal mitochondrial respiration was measured by the attainment of the initial OCR readings and subtraction of the OCR values after treatment with 10 μM antimycin A and 10 μM rotenone (Sigma-Aldrich). Maximal respiration was measured by the subtraction of the nonmitochondrial respiration by the maximum rate measurement after 1 μM carbonyl cyanide 4-(trifluoromethoxy) phenylhydrazone (FCCP) injection. Glycolysis was determined by subtraction of the last rate measurement before glucose injection by the maximum rate measurement before oligomycin injection. Glycolytic capacity was measured by subtraction of the last rate measurement before glucose injection by the maximum rate measurement after oligomycin injection. 2-Deoxyglucose (Sigma-Aldrich; 50 mM) was used to return ECAR to baseline. Experiments were performed in DMEM with no glucose or bicarbonate containing 2 mM glutamine (Sigma-Aldrich).

### RNAseq and qPCR

RNA was extracted using RNAeasy kit (Bio-Rad) and used to perform RNA-seq or to perform qPCR as previously described (Khan et al., 2019). Sequences of primers used are presented in Supplementary Table 1. Libraries preparation, sequencing, and bioinformatics analysis of RNAseq were performed by Novogen (Novogen, Inc, Sacramento CA). Briefly, RNA integrity was assessed with Agilent Bioanalyzer 2100 to select RNA samples with RIN >7.3 to 9.3. Two hundred fifty to 300 base pair insert cDNA libraries, non–strand-specific, were prepared with New England Biolabs (Ipswich, MA) Next Ultra RNA Library Prep and sequenced with Illumina (San Diego, CA) HiSeq PE150 Platform ∼6G/sample Q30 >90%. The reads were mapped to the human reference genome sequence using STAR v2.5 and v2.6.1, with a total mapping rate >90%/sample. For gene expression level analysis and to calculate the fragments per kilobase of transcript per million mapped reads, HTSeq v0.6.1 was used. The differential expression analysis between 2 different groups was done with DESeq2 R package v2_1.6.3. The P values were adjusted using the Benjamini-Hochberg approach for controlling the false discovery rate, adjusted P <.05. TFCat and Cosmic databases were used to annotate the differential expressed gene. The enrichment analysis was done with cluster Profiler R package. The high-throughput sequencing data from this study have been published in GEO with the accession number XXXX (submitted to GEO).

### RNA-seq data download and normalization method

The sample manifest along with the biospecimen sample details sheet and clinical details sheet for all RNA-Seq samples for the TCGA_LIHC (Liver cancer) project were downloaded from The Cancer Genome Atlas (TCGA) Data Portal (https:portal.gdc.cancer.gov) on 18th March 2021. Counts table for all the sample were downloaded using gdc-client 1.6 (https://gdc.cancer.gov/access-data/gdc-data-transfer-tool). The data were normalized as log CPM (Counts Per Million) using edgeR (Robinson et al., 2010), including TMM normalization (Robinson et al., 2010).

### Subcellular fractionation

Fractionation experiments were performed as described before with some modification (Dimauro et al., 2012). Briefly, 1×10^7^ cells were homogenized in a hand-held tight-fitting Teflon pestle homogenizer with 40 strokes in STM buffer (250 mM sucrose, 50 mM Tris–HCl pH 7.4, 5 mM MgCl2, protease and phosphatase inhibitor cocktail) and kept on ice for 30 minutes. The homogenate was centrifuged at 800 X g for 15 mins to separate nuclear fraction as a pellet. The nuclear fraction was washed twice in STM buffer and resuspended in NET buffer (20 mM HEPES pH 7.9, 1.5 mM MgCl2, 0.5 M NaCl, 0.2 mM EDTA, 20% glycerol, 1% Triton-X-100, protease and phosphatase inhibitors). The supernatant was centrifuged at 11000 X g for 10 mins to separate the cytosolic fraction as supernatant and the mitochondrial faction as pellet. The mitochondrial pellet was washed once and resuspended in SOL buffer (50 mM Tris HCl pH 6.8, 1 mM EDTA, 0.5% Triton-X-100, protease and phosphatase inhibitors). All steps were performed on ice and fractions were stored at −80°C.

### Survival analysis of cancer patients with differentially expressed HKDC1

Kaplan-Meier Plotter (KM plotter, http://kmplot.com/analysis/) compiles publicly available data from repositories such as Gene Expression Omnibus (GEO), European Genome-Phenome Archive (EGA), and The Cancer Genome Atlas’ (TCGA). To examine the prognostic value of HKDC1 mRNA expression in HCC, Pan-cancer RNA-seq database was used to evaluate the overall survival of cancer patients (n = 364). The two patient cohorts showing differential gene expression were compared by a Kaplan-Meier survival plot, and the hazard ratio with 95% confidence intervals and logrank P values were calculated.

### TCGA dataset mining

Data from the publicly available TCGA dataset was mined using the websites (https://cistrome.shinyapps.io/timer/) and http://gepia2.cancer-pku.cn/#index.

### Transfection and stable cell line generation

Transient transfection with HK2 and HKDC1 siRNA (Sigma; SASI_Hs02_0035783, SASI_Hs01_0008010) was employed to knockdown (KD) HK2 and HKDC1 protein expression in HepG2 and Hep3B2 cells. Experiments were conducted by plating cells in a 6 well plate for next day transfection when cells where 70-80% confluent using Lipofectamine RNAiMax (Invitrogen; 13778100). A final concentration of (20 pmoL) siRNA was used in culture media without antibiotics overnight after which media was changed to normal growth media and cells were allowed to grow for another 24hrs.

Lentiviruses to overexpress HKDC1-FL (pLV[Exp]-Bsd-EF1A>hHKDC1[NM_025130.4]/HA (VB210314-1076pxg) >10^8^ TU/ml) and HKDC1-TR (pLV[Exp]-Puro-CMV>(Human-HKDC1-TR)/HA (VB200213-1112yyx) >10^8^ TU/ml) were purchased commercially (VectorBuilder). Cells were infected with the 100μL of the virus (in the presence of polybrene; 5 μg/mL) and selected with Blasticidin (10 μg/ml) for 15 days.

For doxycycline inducible shRNA mediated HKDC1 knockdown, Hep3B2 cells were transfected with 4 μg of HKDC1 shRNA construct (Dharmacon; RHS4740-EG80201) using Lipofectamine LTX with Plus Reagent (Invitrogen; A12621). 48 hours post transfection transfected cell were selected with puromycin (0.5 μg/ml) for 3 days. To examine the cells for RFP expression 24-48 hrs post transfection, 1μg/ml doxycycline was used.

### Transmission Electron Microscopy

Cell pellets were fixed in a buffered solution of 2% paraformaldehyde + 2.5 % glutaraldehyde (pH 7.4), washed in 0.1M Sorensen’s sodium phosphate buffer (SPB, pH 7.2) and post-fixed in buffered 1% osmium tetroxide for 1 hour. After several buffer washes, samples were dehydrated in an ascending concentration of ethanol leading to 100% absolute ethanol, followed by 2 changes in propylene oxide (PO) transition fluid. Specimens were infiltrated overnight in a 1:1 mixture of PO and LX-112 epoxy resin, and 2 hours in 100% pure LX-112 resin, and then placed in a 60 °C oven to polymerize (3 days). Ultra-thin sections (∼75 nm) were cut (using a Leica Ultracut UCT model ultramicrotome), collected onto 200-mesh copper grids and contrasted with uranyl acetate and Reynolds’ lead citrate stains, respectively. Specimen were examined via JEOL JEM-1400F transmission electron microscope, operating at 80 kV. Digital micrographs were acquired using an AMT Biosprint 12M-B CCD Camera and AMT software (Version 7.01).

### Statistical analysis

Values are represented as means ± standard errors of the mean (SEM). Data were analyzed ether by student’s *t*-test or 1-way/2-way ANOVA followed by a Tukey or Bonferroni post-hoc test when applicable. Analysis of RNAseq data and enrichment analysis of DEG was performed by Novogen, Inc. Differentially regulated metabolites and enrichment analysis of metabolomics was performed with Metaboanalyst software. The statistical analyses were performed using GraphPad Prism 8 (GraphPad Software, La Jolla, CA). p-values less than 0.05 were considered significant.

## Results

### HKDC1 is overexpressed in HCC

Since our discovery of HKDC1 as a novel hexokinase and the initial observation that it is overexpressed in HCC (Pusec et al., 2019, Zhang et al., 2016), we have further mined data from the TCGA datasets observing that HKDC1 is significantly upregulated in a wide variety of human cancer types (Suppl Fig 1A). Exploring this further, we observed that HKDC1 has high expression in tumors of HCC patients (Fig 1A) and that HKDC1 expression was significantly higher in different stages (I to III) of HCC in these patients (Fig 1B). Further, HKDC1 expression was associated with lower survival in these patients (Fig 1C). Interestingly, further exploration shows that HKDC1 is the hexokinase isoform most significantly upregulated in HCC (Supplementary Fig 1B). Importantly, in HCC, it has been previously shown that there is a major metabolic reprogramming, in part, due to a shift from the primary hepatic hexokinase (HK) isoform, glucokinase (GCK) to HK2 (DeWaal et al., 2018) indicating that HK2 plays a key role in HCC. Since HK2 knockdown does not completely block the development of HCC (DeWaal et al., 2018), these data indicate that other HKs may also have a role and our findings suggest that HKDC1 may play this role.

Non-alcoholic fatty liver disease (NAFLD) is a known independent risk factor for HCC (Nagaoki et al., 2021, Orci et al., 2021), and we have previously shown that higher liver HKDC1 expression correlates with hepatic steatosis (Pusec et al., 2019). Therefore, we used a mouse model to explore HKDC1 expression in mice fed a NASH diet (high-fat, cholesterol and sucrose diet (HF-HC-HSD) as compared to mice fed a nutrient-matched low fat, low cholesterol and low sucrose diet as controls (LF-LC-LSD; Con) for 34 weeks, where HCC develops in a subset of mice on this diet. In the NASH diet group, 100% of mice developed NASH and 15% of that group developed HCC in addition to NASH (NASH+HCC). HKDC1 expression (mRNA and protein), as expected, was significantly overexpressed (» 10-fold) in the livers of the NASH diet fed mice as compared to controls (Fig 1D; left panel). More intriguingly, mice that developed HCC along with NASH had » 100-fold higher HKDC1 mRNA expression in the liver tissue (non-tumor regions) than controls (Fig 1D; left panel). This increase in HKDC1 mRNA expression was also evident in enhanced HKDC1 protein expression (Fig 1D; right panel). This dramatic progressive increase in hepatic HKDC1 expression from NASH to HCC suggests an association of HKDC1 in the progression of liver disease to HCC but does not demonstrate a causative role.

### HKDC1 expression is essential for HCC growth and proliferation

Since, HKDC1 has very low expression in the adult liver (Khan et al., 2018, Pusec et al., 2019), and its expression is upregulated in HCC (Zhang et al., 2016, Cerami et al., 2012), we hypothesized that the upregulation of HKDC1 expression in HCC promotes proliferation in HCC. Testing this hypothesis, we used AML12 cells, a non-cancerous hepatocyte cell line, which has minimal expression of HKDC1, and tested for the effect of stable overexpression of HKDC1 (Supp Fig 1C). We found that HKDC1 overexpression resulted in enhancement of both proliferation (Supp Fig 1D) and survival capacity (Supp Fig 1E). To further investigate the effect of HKDC1 expression on HCC survival, we selected 5 different HCC cell lines with variable HKDC1 expression (Suppl Fig 2A) and downregulated HKDC1 expression using siRNAs against HKDC1 (Suppl Fig 2B). HKDC1 downregulation inhibited cell survival *in vitro* in the classical colony forming assay (Supplementary Fig 2C) regardless of the endogenous expression of HKDC1 in these cell lines. These data indicate that HKDC1 is required for survival and proliferation in these cell lines. To further explore this phenotype, we developed a HKDC1 knockout (KO) cell line by using Crispr-Cas9 technology in HepG2 cells (Supplementary Fig 2D-E) and used these modified HKDC1-KO cells to perform an *in vitro* viability and proliferation assay. These assays showed that knock down of HKDC1 resulted in diminished proliferation and survival (Fig 2A-B) as well as reduced expression of proliferation marker, Ki67 (Fig 2C). This suggests a potential role of HKDC1 in HCC proliferation.

**Fig 2.**
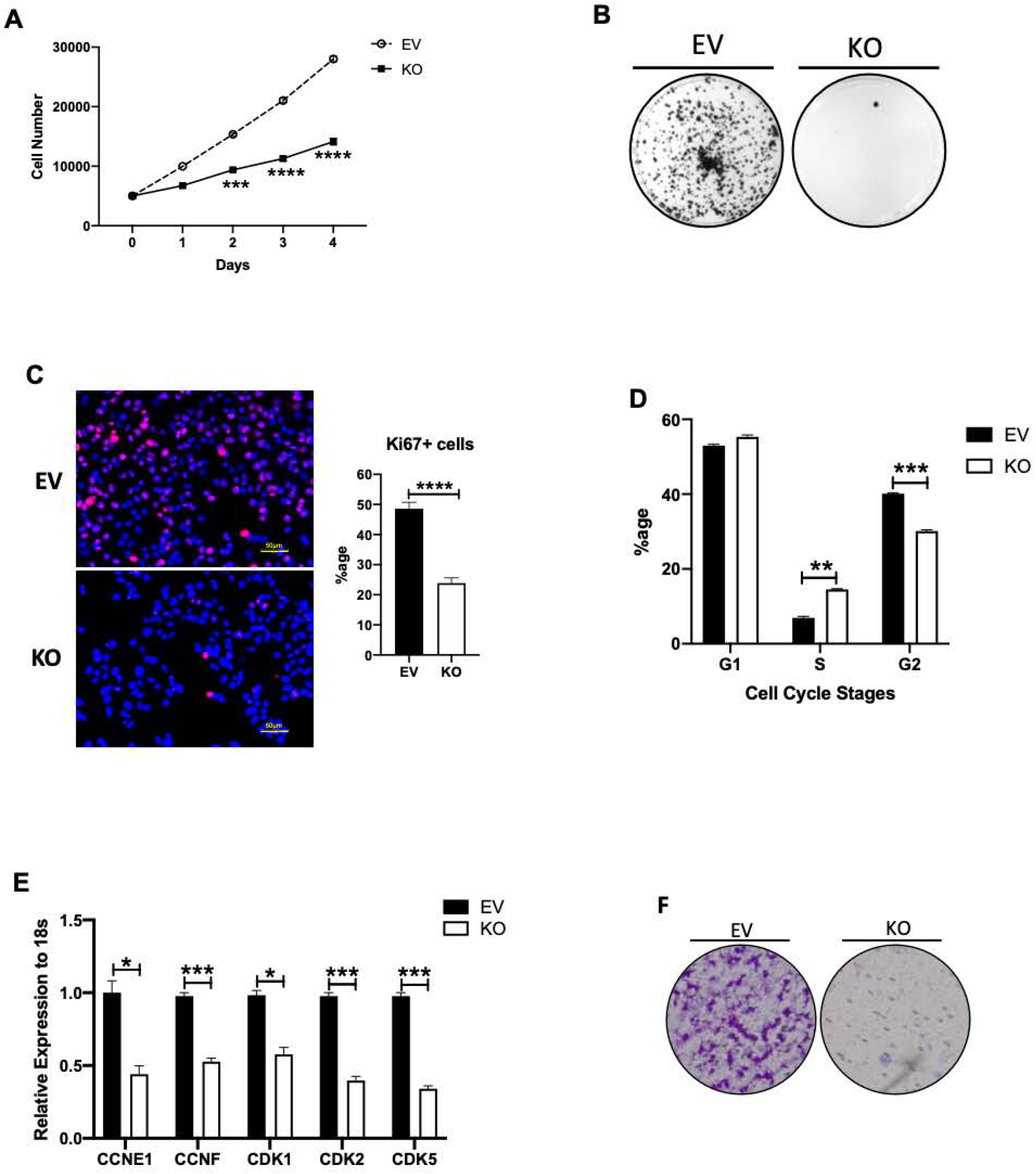
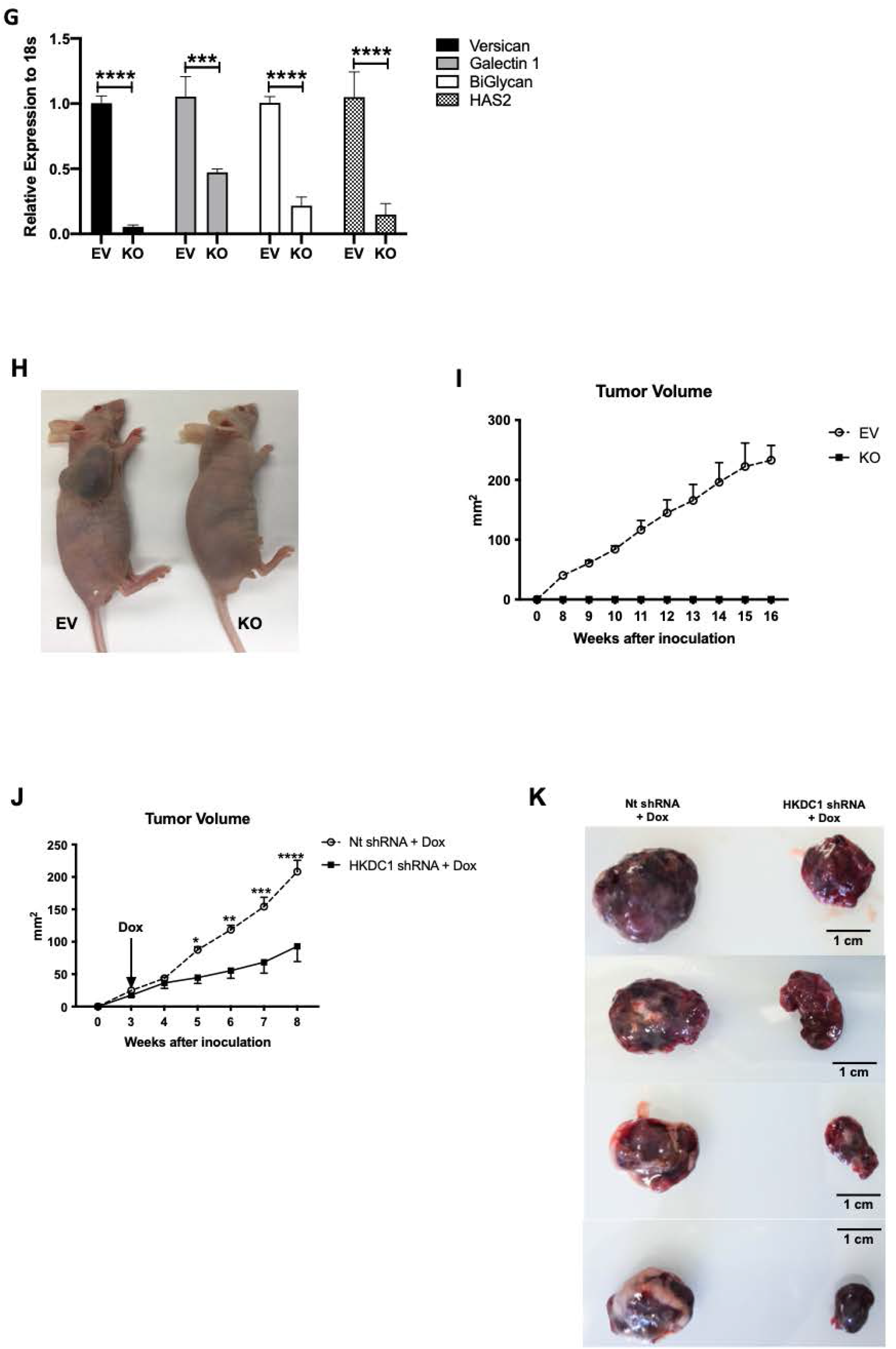
HKDC1 is essential for HCC progression and survival. EV and HKDC1-KO cells (KO) were used for **A)** cell proliferation assay, **B)** colony forming assay, **C)** immunostaining for Ki67 proliferative marker (pink), nuclei are stained with DAPI (blue), **D)** cell cycle analysis with propidium iodide, **E)** mRNA expression by qPCR for cyclins and cyclin dependent kinases vs KO cells, **F)** transwell migration assay to assess invasiveness and **G)** mRNA expression by qPCR for proteoglycan synthesis genes. **H)** *in vivo* tumor growth was assessed where 1×10^6^ EV or KO cells were inoculated into 4-6 weeks old male Nu/J mice (n=5), images were taken at endpoint (16 weeks post inoculation). **I)** Tumor size was measured weekly till 16 weeks after appearance of tumor with a vernier caliper tumor weight till the end of the study. **J)** Hep3B2 cells were transfected with shHKDC1 or (non-target) ntshRNA and transfected cells were selected with appropriate antibiotics, 1X10^6^ cells were inoculated into mice (n=4). When tumors were visible, mice were given doxycycline (in diet) for 7 days to activate shRNAs. Tumor growth was measured weekly till 8 weeks after appearance of tumor with a vernier caliper. **K)** Images of tumors at endpoint. All cell line experiments (**A-G**) were performed 2-3 times with 3-5 replicates per experiment. Values are mean ± SEM; *p < 0.05; **p < 0.01; ***p < 0.001; ****p < 0.0001 by Student’s t-test or 2-way ANOVA (for 2J).

To investigate the reduction in proliferation of HCC cells lacking HKDC1 (KO), we analyzed gene expression in HKDC1-KO cells as compared to control cells (carrying the empty vector; EV) by RNA-seq. This showed that there were significant genome wide changes with the HKDC1-KO cells with 3000 genes significantly upregulated or downregulated in HKDC1-KO cells compared to EV cells (Supplementary Fig 2F). Moreover, there were 449 genes that were uniquely expressed in HKDC1-KO cells and 1199 genes absent that would normally be expressed which would normally be expressed in HepG2 cells (Supplementary Fig 2G). Of the top 30 significantly downregulated gene ontology (GO) terms in the gene ontology analysis, 29 GO terms were related to cell division machinery (Supplementary Fig 2H) consistent with the observed reduction in proliferation rates in HKDC1-KO cells. To analyze this further, we performed cell-cycle analysis on the cells and the data indicates that in HKDC1-KO cells, there is a significant increase in cells in synthesis (S) phase (in which cells synthesize DNA) and a concomitant significant decrease in G2 phase (Fig 2D). An increase in cell populations in S phase could be due to failure in DNA synthesis machinery to progress through this phase. Cyclins and cyclin dependent kinases (CDKs) act as tumor suppressors by controlling cell growth and inducing the death of damaged cells (Lim and Kaldis, 2013, Wenzel and Singh, 2018). Genetic mutations, as in cancer, causes the malfunction of one or more of these regulatory proteins at cell cycle checkpoints resulting in uncontrolled cellular replication, leading to carcinogenesis, or tumor development (Lim and Kaldis, 2013, Wenzel and Singh, 2018). Our RNA-seq data showed significant downregulation for several cyclins and CDKs, which we validated by qPCR analysis (Fig 2E). Thus, our data indicate that HKDC1 ablation results in the downregulation of cyclins and CDKs, which are key factors in cell cycle progression and proliferation.

Cell migration abilities are involved in pathological processes such as cancer metastasis (Trepat et al., 2012). Cancer cells must migrate and invade through the extracellular matrix (ECM) to metastasize. Testing whether HKDC1 has a role in cell migration, we used a classical trans-well assay to assess the effect of HKDC1 ablation. In this assay HKDC1-KO HepG2 cells lost the ability to migrate and invade properties when compared to EV cells (Fig 2F). Further, since the property of invasiveness depends on the synthesis of proteoglycans (Wight et al., 1992, Snigireva et al., 2019), we searched for the expression levels of important genes involved in proteoglycan synthesis in our RNA-seq data and found that they were significantly downregulated when HKDC1 was ablated in HepG2 cells, which we confirmed by qPCR (Fig 2G).

Next using an *in vivo* xenograft model where we injected the modified HepG2 cell lines in immunocompromised mice (Nu/J mice), we observed that none of the immunocompromised mice injected HKDC1-KO cells formed tumors, whereas 100% of mice injected with EV cells developed tumors (Fig 2H-I, and Supplementary Fig 2I). To further investigate this, we created an inducible shRNA mediated knockdown of HKDC1 in a different HCC cell line that has high HKDC1 expression (Hep3B2 cells) and performed xenografts experiments with these cell and control cells carrying a scrambled shRNA (Supplementary Fig 2J). Our data shows that although tumors form in both cell lines (containing scramble and shHKDC1), when the shRNA expression was induced to knockdown HKDC1 using a doxycycline-diet, the tumors in shHKDC1 group of mice exhibited reduced growth as compared to cells carrying scrambled shRNA (Nt shRNA) (Fig 2J-K; Supplementary Fig 2K). To further test if this finding can be replicated in another model of HCC, we used the classical chemical based (diethylnitrosamine; DEN) induction of hepatic carcinogenesis. For this experiment, we generated a mouse model by crossing our previously described HKDC1 floxed (HKDC1^f/f^) mice (Khan et al., 2019, Pusec et al., 2019) with Albumin-Cre mice to generate liver specific HKDC1 knockout mouse (HKDC1-LKO). DEN was injected at 14 days and the animals were observed for 40 weeks (Supplementary Fig 3A-C). As a proof of concept, we re-expressed full-length human HKDC1 in mice from both HKDC1^f/f^ and HKDC1-LKO groups (Supplementary Fig 3A). At the terminal time point (40 weeks), we found that the HKDC1^f/f^ (controls) had larger livers than HKDC1-LKO mice (Fig 3A-B; Supplementary Fig 3D) which had both fewer and smaller tumors as compared to the control animals (Fig 3A-B; Supplementary Fig 3D). More importantly, mice from both control and HKDC1-LKO groups where HKDC1 was overexpressed (by HKDC1 AAV) had both larger livers and more tumors (Fig 3A-B; Supplementary Fig 3D). Further, histological analysis revealed that livers from HKDC1-LKO mice with AAV treatment had the highest number of BrdU and Ki67 positive hepatocytes compared to all groups while HKDC1-LKO had the least number of positive cells compared to any other group (Fig 3C-D) Thus, our data from both *in vitro* and *in vivo* HCC models indicates that HKDC1 regulates several important cellular processes and establishes its role in the proliferation and survival of HCC.

**Fig 3.**
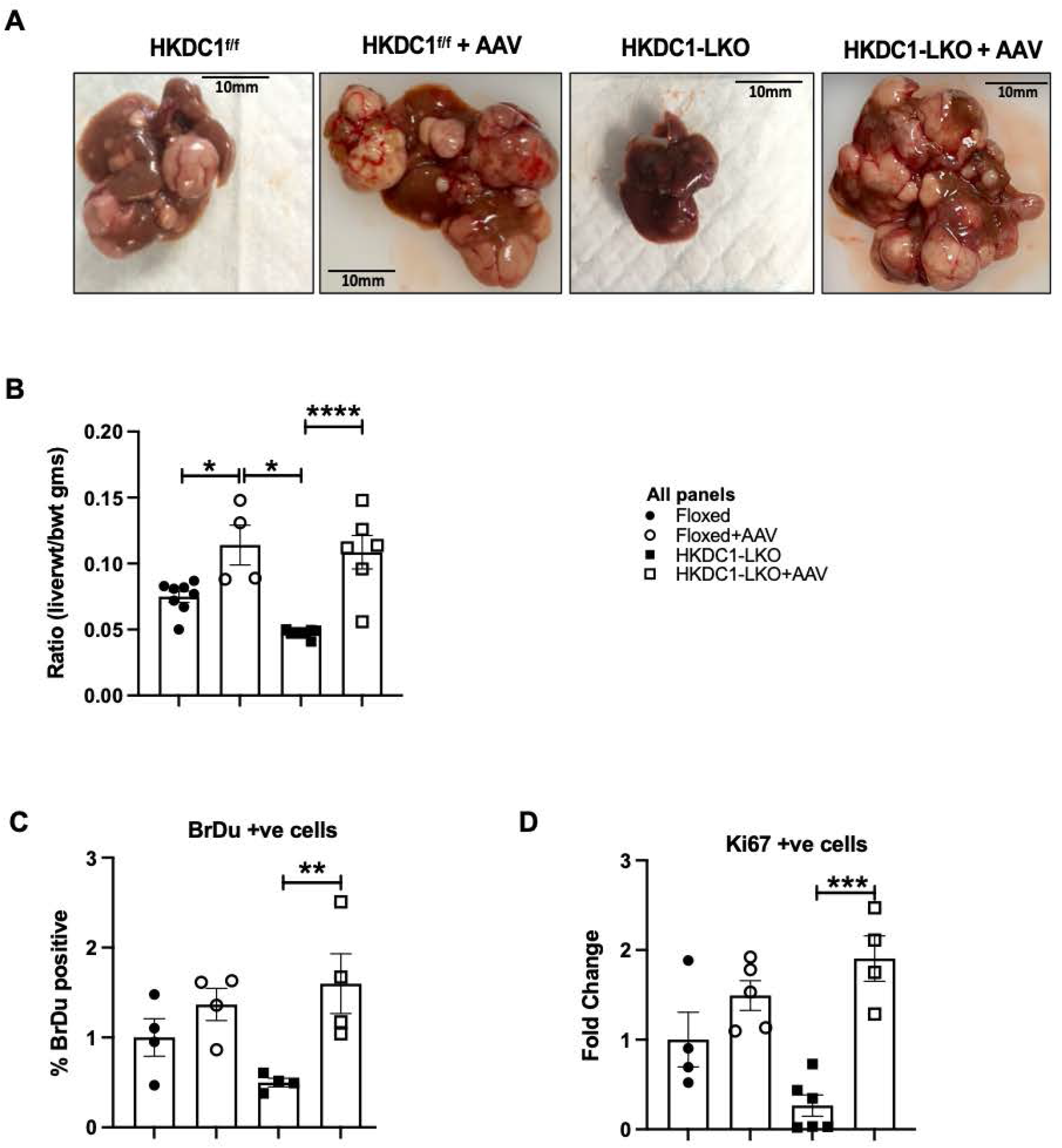
HKDC1 ablation impairs HCC progression in an *in vivo* HCC model. Two-week-old HKDC1^f/f^ and HKDC1-LKO male mice were injected with DEN (25 mg/kg). When mice were 8 weeks old, both groups were further divided into groups where one group received AAV expressing human HKDC1 (HKDC1^f/f^+AAV and HKDC1-LKO+AAV) and the AAV expressing null vector was used as the control with the two other groups (HKDC1^f/f^ and HKDC1-LKO), with N=3-7 per group. Ten months after DEN injection, mice were sacrificed and **A)** images of the livers are shown. **B)** Livers were weighed and ratio to body weight was calculated. Liver sections were fixed, and immunohistochemistry was performed to show **C)** BrdU and **D)** Ki67 positive hepatocytes. Regions of tumor in the histology slides were identified and omitted from analysis. Values are mean ± SEM; *p < 0.05; **p < 0.01; ***p < 0.001; ****p < 0.0001 by 2-way ANOVA.

### HKDC1 ablation disrupts glucose metabolism

Human HCC cells generally do not express HK1 but have enhanced expression of HK2, the knockdown of which results in a reduction in cell proliferation and reduction in glycolysis (DeWaal et al., 2018). Analysis of HK1 and HK2 expression has revealed a reciprocal pattern of expression of these HKs in many normal and cancerous cells, suggesting that expression is regulated to maintain total HK activity or G-6-P quantity (Jiang et al., 2021, Garcia et al., 2019, Patra et al., 2013). Since HKDC1 is a putative hexokinase gene, we examined whether ablating *HKDC1* gene expression would affect other HKs, including their expression, cellular localization and total cellular hexokinase activity. To investigate HK expression, we assessed the mRNA and protein levels of HK1, HK2 and GCK in the HKDC1-KO cells as compared to EV and found that ablation of HKDC1 had no effect on the mRNA or protein levels of the other HKs in HepG2 cells (Supplementary Fig 4A-B).

Next, we assessed the impact of HKDC1 ablation on cellular hexokinase activity. Under varying glucose concentrations, total hexokinase activity in the HKDC1-KO cells was not significantly different from EV cells (Fig 4A). Since HK2 is the other predominant HK isoform in HCC and it has been previously shown that knockdown of HK2 impacts glycolysis (DeWaal et al., 2018), we used siRNAs against either HK2 or HKDC1 to determine the relative impacts of these HKs on glycolysis in HepG2 cells. Our data shows that when HK2 was knocked down (KD) in HepG2 cells, there was ∼ 40% reduction in cellular HK activity (Fig 4B; Supplementary Fig 4C). On the contrary, when HKDC1 was knocked down in, we did not observe any effect on HK activity (Fig 4B; Supplementary Fig 4C). We then performed Seahorse assays in our HepG2 cells (Crispr-KO) and two other HCC cells lines, Hep3B2 and SNU-475 (using siRNAs to knock down HKDC1 expression). The data shows that there is no change in glycolysis in HepG2 and SNU-475 cells with HKDC1 knockdown while there was an increase in glycolysis in Hep3B2 cells. Further there was no change in glycolytic capacity in all the cell lines upon HKDC1 KO or KD (Fig 4C). This data is consistent with HKDC1 ablation not affecting HK activity, confirming that HKDC1 is a non-canonical HK in HCC. To further study this observation, we used AML12 cells with and without HKDC1 overexpression and found that HKDC1-OE had no effect on glycolysis or glycolytic capacity (Supplementary Figure 4D). Overall, these data are consistent with our earlier studies which show that HKDC1 is a poorly functioning HK (with low Km), as compared to other HKs (Pusec et al., 2019). Taken together our data suggest that while both HK2 and HKDC1 are upregulated in HCC, HK2 has a direct role in enhancing glycolysis (DeWaal et al., 2018), while HKDC1 does not mediate this role.

**Fig 4.**
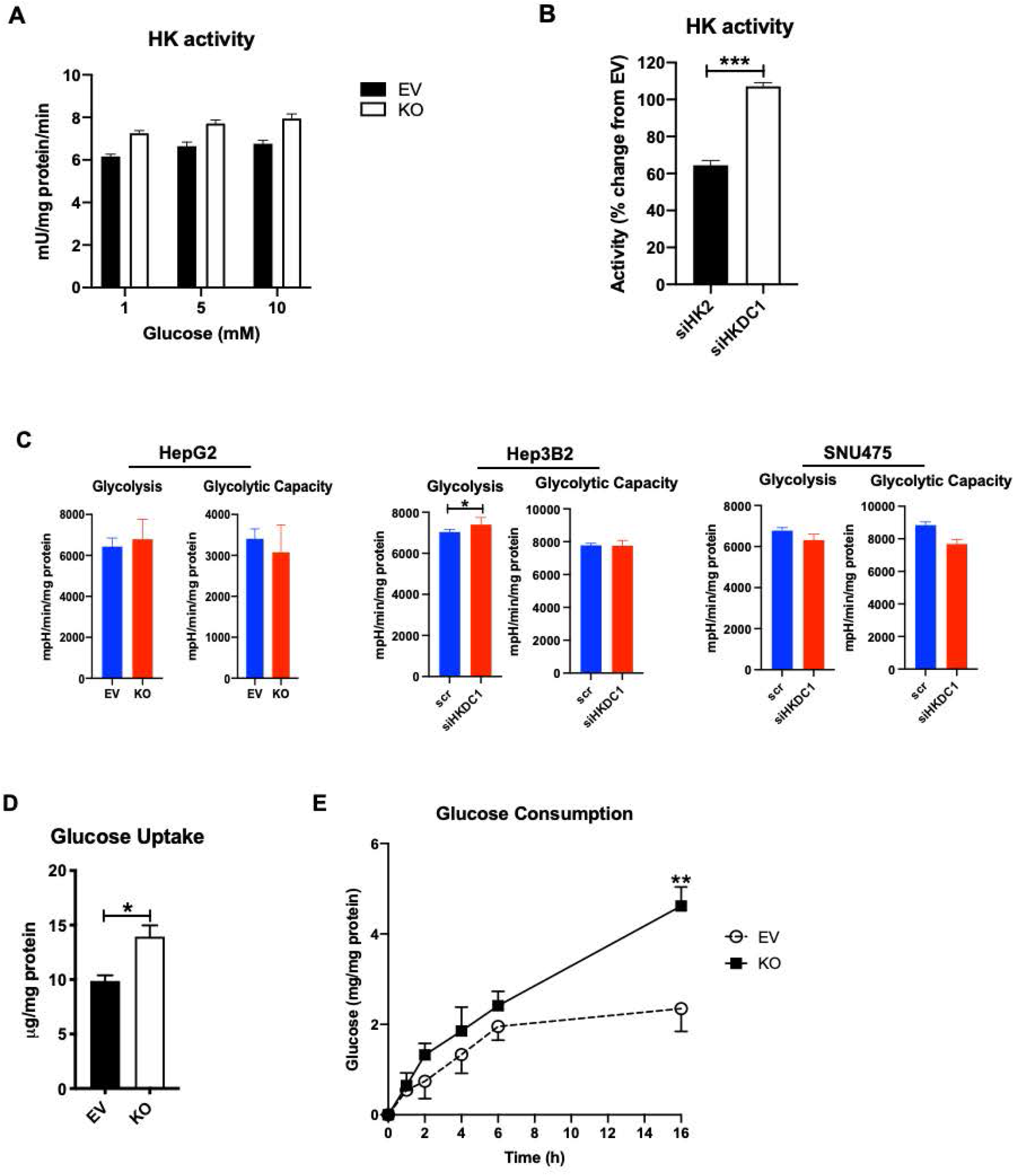
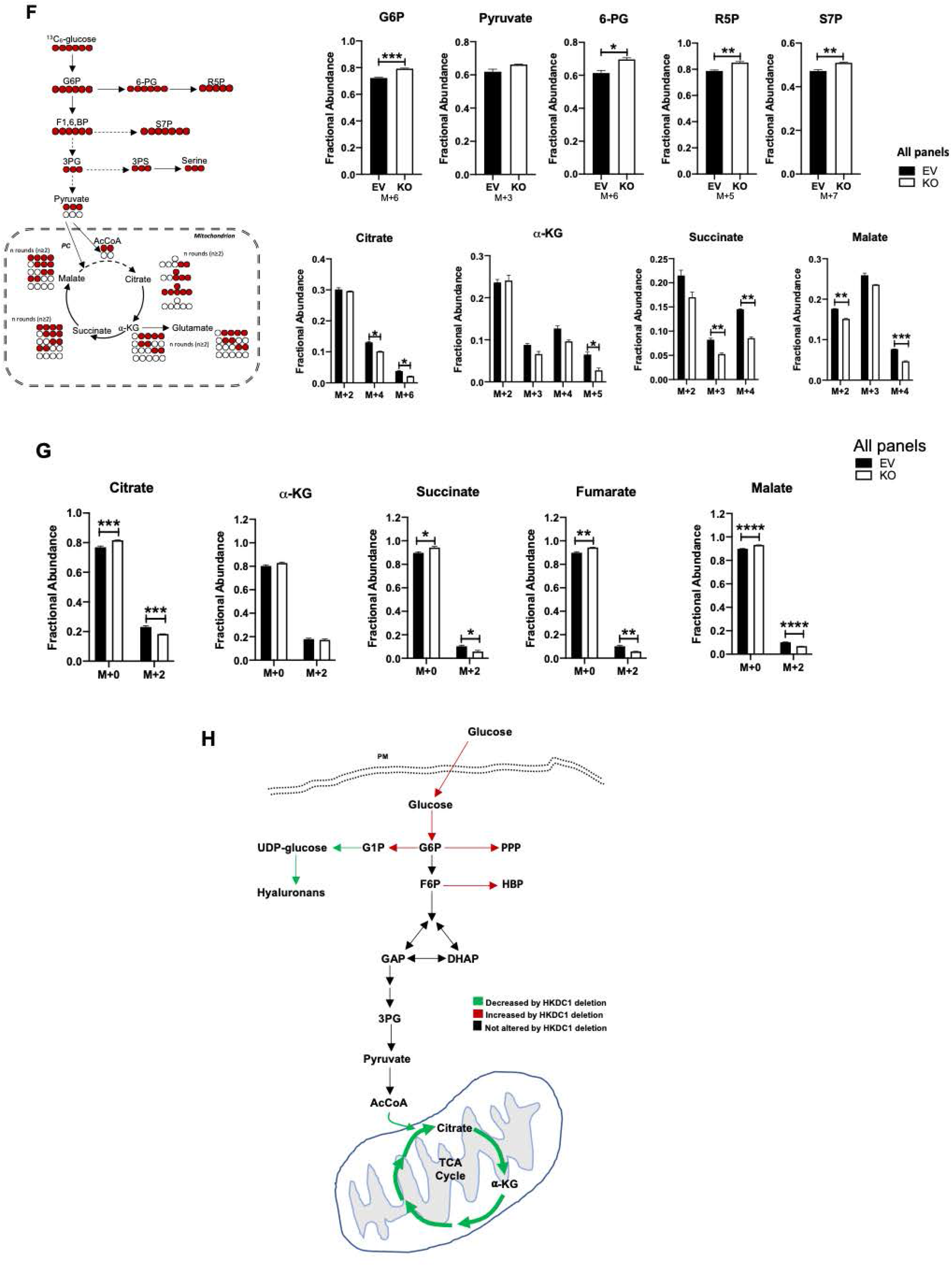
HKDC1-KO impairs glucose metabolism. **A)** Hexokinase activity in EV and KO HepG2 cells. **B)** HepG2 cells were treated with siRNA against either HK2 or HKDC1 for 24h, cells were lysed, and hexokinase activity was assayed. **C)** Seahorse metabolic analysis (ECAR) of EV and HKDC1-KO cells (left panel) and siRNA mediated HKDC1 knockdown (siHKDC1) in Hep3B2 and SNU475 cells (center and right panels). **D)** 2-NBDG (2-(N-(7-Nitrobenz-2-oxa-1,3-diazol-4-yl)Amino)-2-Deoxyglucose) fluorescent analog of glucose was used to assess glucose uptake in EV and KO HepG2 cells. **E)** In EV and KO HepG2 cells, glucose consumption was assessed by measuring glucose concentration in media aliquots taken at designated time periods which was subtracted from initial glucose concentration of media, obtaining glucose being consumed by the cells. **F)** Mass isotopomer analysis of glycolytic and TCA cycle metabolites in cells cultured with 5.5 mM of [U-^13^C_6_] glucose and unlabeled glutamine (n=3, independent biological replicates) for 4h. **G)** Mass isotopomer analysis of TCA cycle metabolites in cells cultured with 2 mM of [U-^13^C_5_] glutamine and unlabeled glucose (n=3, independent biological replicates) for 4h. **H)** Schematic summarizing the changes in glucose flux from Table1 and 4E-F). Exp 4A-D were performed 2-3 independent times, with three replicates per individual experiment. Values are ± SEM; *p < 0.05; **p < 0.01; ***p < 0.001; ****p < 0.0001 by Student’s t-test (for 4A-C) or 2 way ANOVA (for 4D-F).

Although HKDC1 has been shown by us previously (Pusec et al., 2019) and here to not be a major driver of cellular hexokinase activity, we next examined if HKDC1 impacts other aspects of glucose metabolism. First, we observed that HKDC1 ablation significantly increases glucose uptake as compared to EV cells (Fig 4D). While GLUT2 is the predominant form in liver cells, GLUT1 is the major form of glucose transporter in HCC (Kim et al., 2017). To assess if the observed increase in glucose uptake could be due to enhanced GLUT expression, we studied the expression of glucose transporters. We observed that GLUT1 levels remain unchanged while surprisingly GLUT2 levels were reduced at both the mRNA and protein levels with HKDC1 ablation (Supplementary Fig 4E-F). Interestingly, the levels of GLUT4, which is known to be activated by the PI3K-Akt pathway (Hong et al., 2016), was significantly upregulated (Supplementary Fig 4E-F). Concomitant with the increased glucose uptake, we found that HKDC1-KO cells consumed significantly more glucose over time as compared to EV cells in an assay where the disappearance of glucose (consumption) from the media was measured (Fig 4E).

Increased glucose consumption is a part of the classical “Warburg” effect (Liberti and Locasale, 2016, Vaupel et al., 2019), where cancer cells have a preference of taking in glucose at much higher rates than normal cells. However, for successful carcinogenesis to take place, a more intricate balance in nutrient use needs to be achieved rather than exorbitant glucose consumption. We suggest that upon HKDC1 ablation, cancer cell metabolism is impacted, resulting in energetic stress and increased glucose consumption as a way of survival. To examine changes in nutrient use more closely, we next investigated the effect of HKDC1 ablation on glucose flux by performing a steady-state metabolomics, focusing on metabolites related to glucose metabolism. Our data shows that there were increased glucose-6-phosphate levels in HKDC1-KO cells which could be a result of the increased glucose uptake/consumption (Table 1). However, we did not find a similar increase in glycolytic metabolites (Table 1). Interestingly, we did find an increase in the levels of metabolites involved in the PPP and HBP shunts of the glycolysis (Table 1). As we observed earlier, an enhanced glucose consumption in HKDC1-KO cells did not result in more pyruvate entering the TCA cycle and therefore, the increase in PPP/HBP metabolites may be from enhanced glucose consumption. To take a step further in our investigation we performed a targeted metabolomics using U-^13^C_6_ labeled glucose to determine how the excess glucose consumed by HKDC1-KO cells was utilized. These data showed that labeled carbons from glucose are converted to glucose-6-phosphate (Fig 4F; upper panel) but they do not accumulate in the glycolytic intermediates, but rather cause an increase in PPP and HBP metabolites (Fig 4F; upper panel). Moreover, there is a significant decrease in labeled glucose carbons entering the TCA cycle (Fig 4F; lower panel). Since anaplerosis feeds metabolites to the TCA cycle (Owen et al., 2002) in addition to glucose we performed a U-^13^C_5_ labeled glutamine targeted metabolomics, and the data shows that indeed on HKDC1 ablation there is a significant decrease in labelled TCA cycle metabolites upon HKDC1-KO (Fig 4G; all panels). As summarized in Fig 4H, cells lacking HKDC1 have enhanced glucose consumption fueling the PPP and HBP pathways but decreased TCA cycle flux.

**Table 1:**
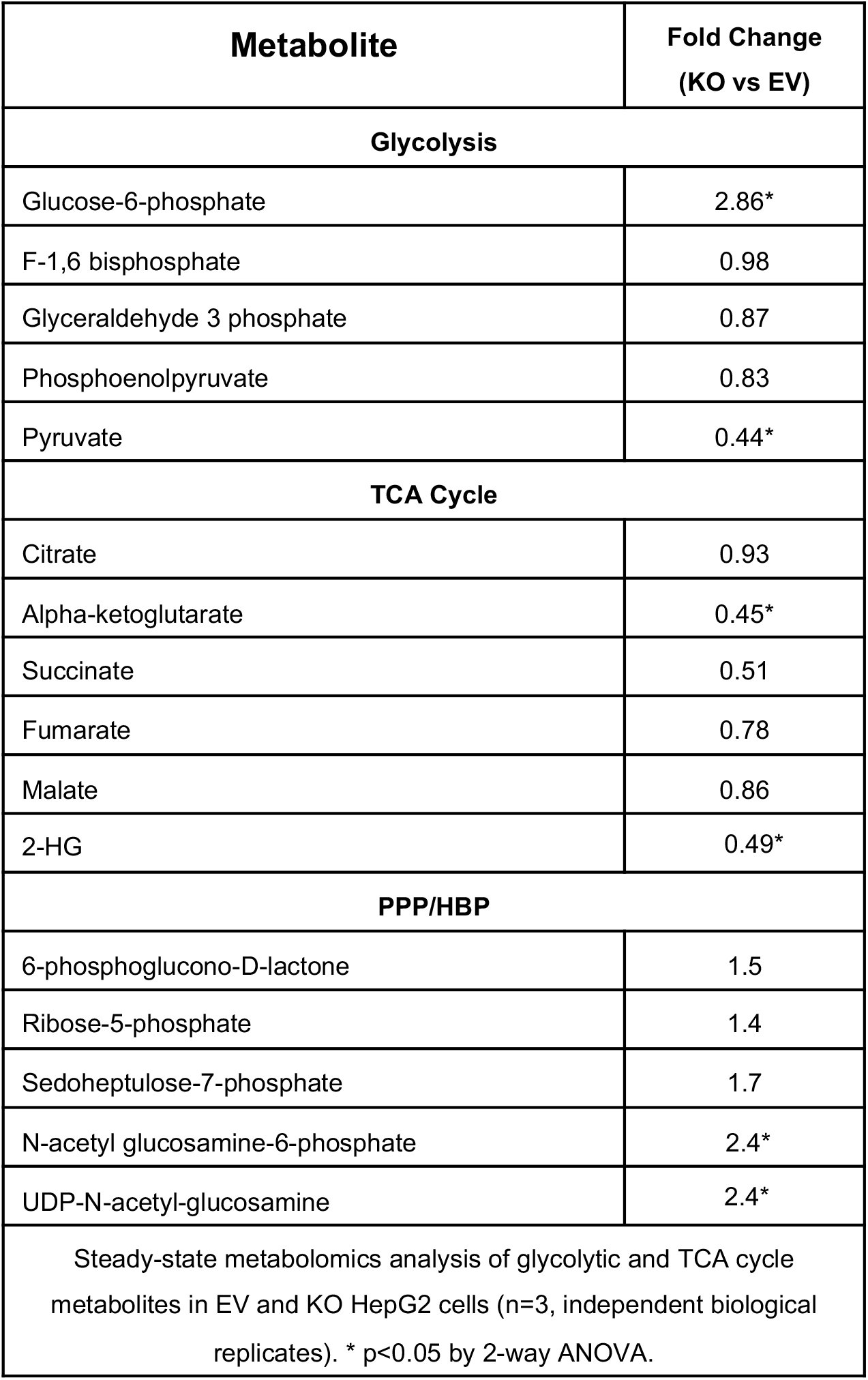
Fold change in metabolite levels upon HKDC1-KO in HepG2 cells.

### HKDC1 is essential for mitochondrial function

Among the HKs, both HK1 and HK2 which interact with the mitochondria have been shown to play critical roles in promoting cell proliferation and survival in breast, colon and prostate cancers, cervical carcinoma, gastric adenoma, glioma and lymphoma (Pastorino et al., 2002, Robey and Hay, 2009, McDonald et al., 2019, Robey and Hay, 2006). As we have previously shown that HKDC1 localizes to the mitochondria and binds with mitochondrial proteins like VDAC (Pusec et al., 2019), we performed a subcellular fractionation experiment in our modulated HepG2 cells, which showed that HKDC1 was predominantly in the membrane fraction with a lesser amount in the cytosol (Fig 5A; right panel). In contrast, HK2, which has been shown to bind to the mitochondria in some cancer cell lines (Majewski et al., 2004, Pastorino et al., 2002), it was more found to be in the cytosol than membrane fraction (Fig 5A; right panel). Interestingly, in HKDC1-KO cells we didn’t see more HK2 in the mitochondrial fraction which suggests that HKDC1 does not compete with HK2 for mitochondrial binding at least in HepG2 cells (Fig 5A; left panel). We confirmed this observation by examining two other HCC cell lines (Hep3B2 and Huh7) and found that in both cell lines, HKDC1 is more at the mitochondria than in the cytosol fraction while the opposite is true for HK2 (Fig 5B; Supplementary Fig 5A).

**Fig 5.**
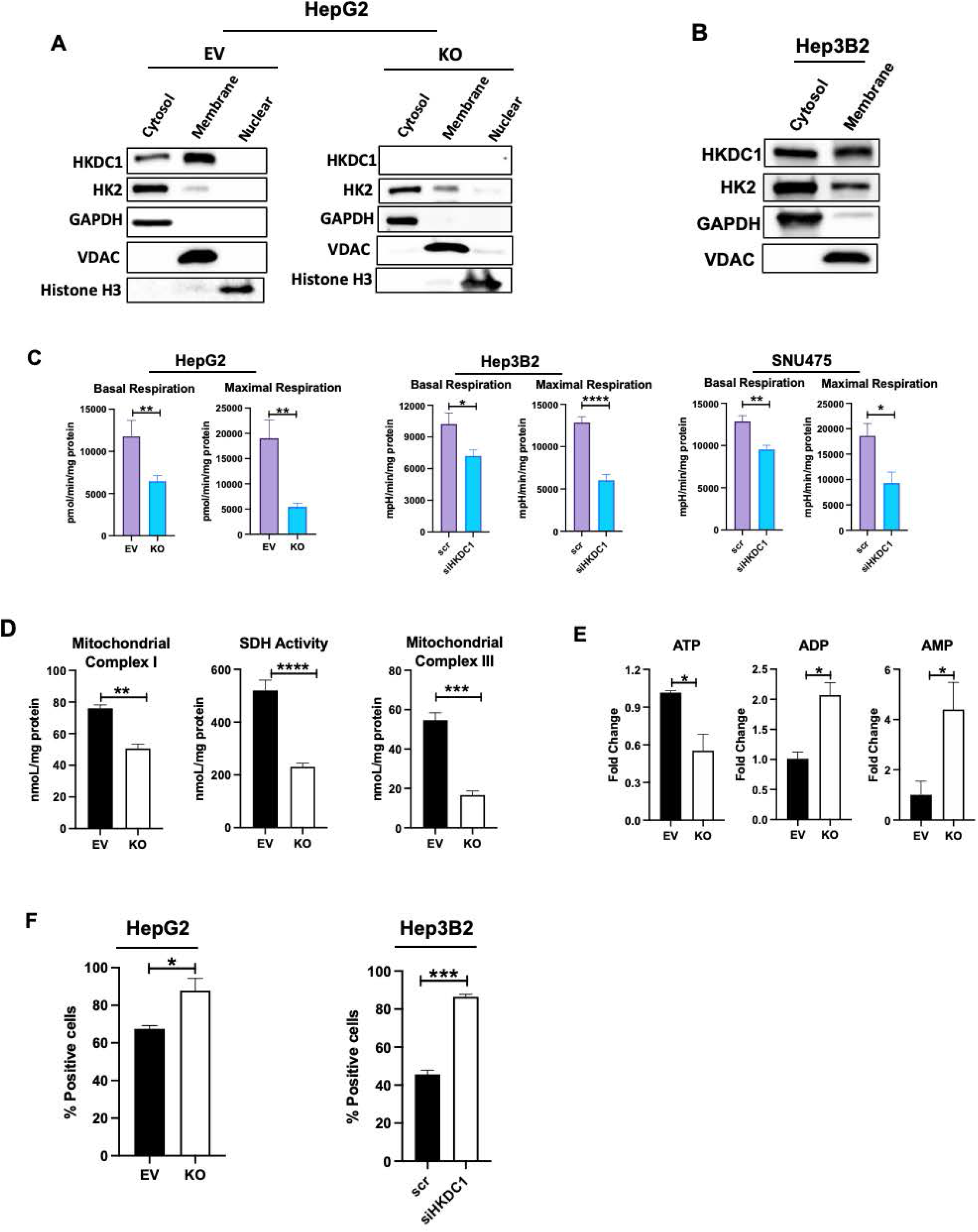
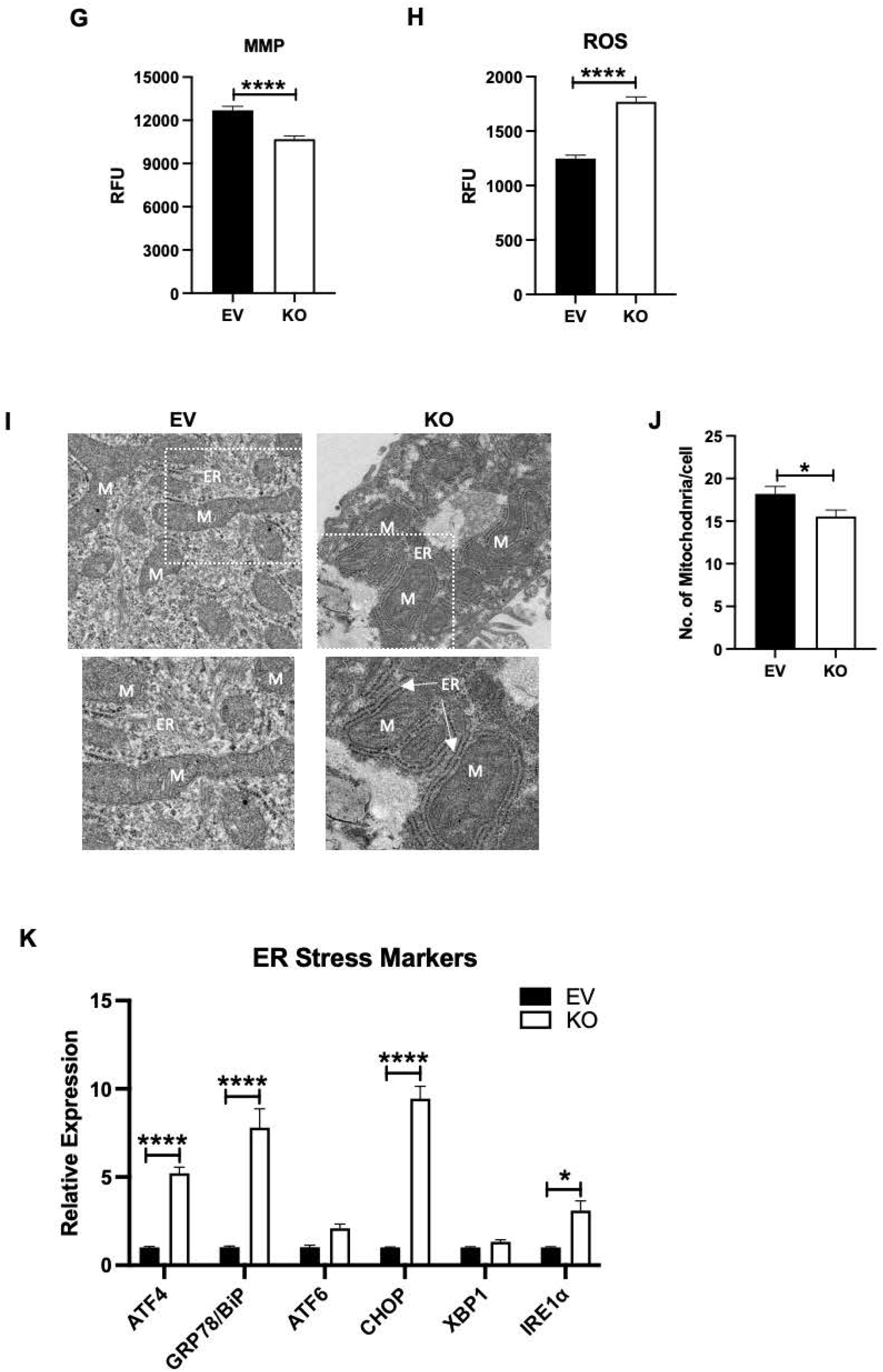
HKDC1 is essential for mitochondrial function in HCC. Cell fractionation experiment showing cytosolic, membrane and nuclear fractions in **A)** EV and KO HepG2 cells and **B)** Hep3B2 cells. **C)** Seahorse metabolic analysis (OCR) of EV and HKDC1-KO cells and siRNA mediated HKDC1 knockdown (siHKDC1) in Hep3B2 and SNU475 cells (center and right panels). **D)** Activity assay for mitochondrial complex I (left panel), SDH (center panel) and complex III (right panel) of EV and HKDC1-KO cells. **E)** Relative levels in fold change (to EV) of ATP, ADP and AMP values from steady-state metabolomics (n=3). **F)** Mitochondrial Ca^2+^ levels were assessed by Rhod-2AM fluorescence using flow cytometry of EV/HKDC1-KO cells (left panel) and Hep3B2 cells treated with siRNA against HKDC1 for 24h (right panel). 50,000 cells were assessed for each sample, and data was plotted on bar graphs with statistics (n=3) for each cell line **G)** Mitochondrial membrane potential was measured by TMRE fluorescence. **H)** Intracellular ROS levels are shown. **I)** HepG2 expressing empty vector (EV) or HKDC1 knockout (KO) cells were processed for TEM. 20-25 images were taken for each cell at different magnifications. Images shown here were taken at 4000X. Inset (shown in white) was enlarged (below) to show mitochondria and ER (M=mitochondria, ER=endoplasmic reticulum). **J)** Mitochondria were counted for each cell (20-27 cells per sample) by Image J software. All cell line experiments were performed 2-3 independent times with 3-8 replicates per experiments. **K)** qPCR analysis in EV and KO cells to assess mRNA levels of ER stress markers. Values are ± SEM; *p < 0.05; **p < 0.01; ***p < 0.001; ****p < 0.0001 by Student’s t-test (for 5B, E-G, I) or 2 way ANOVA (for 5C,J).

Mitochondrial function is essential for carcinogenesis as shown by numerous studies (Hsu et al., 2016, Luo et al., 2020, Srinivasan et al., 2017). Exploring our RNA-seq data shows that of the top 20 significantly downregulated GO terms (cellular component) upon HKDC1-KO included several belonging to the mitochondria (Supplementary Fig 5B); thus, we reasoned that HKDC1 might be required for optimal mitochondrial function in HCC. To investigate this hypothesis, we performed Seahorse analysis in our HepG2 cells (HKDC1-KO by Crispr-Cas9) and two other HCC cells lines, Hep3B2 and SNU-475 (using siRNAs to KD HKDC1) observing that both basal and maximal respiration was significantly reduced in HKDC1-KO/KD cells suggesting that HKDC1 is important for mitochondrial function (Fig 5C). To further explore, we measured activities of mitochondrial complex I, SDH and III and observed that HKDC1 ablation significantly reduces the activities of all three complexes (Fig 5D). Cellular energy in the form of ATP mainly comes from mitochondrial activity even in cancer cells and enhanced glucose consumption observed in HKDC1-KO cells cannot provide this energy as indicated by the significantly reduced levels of ATP (Fig 5E).

Calcium influx from the ER is essential for mitochondrial metabolism, where a transient increase in Ca^2+^ level activates matrix enzymes and stimulates oxidative phosphorylation, but sustained exposure to high Ca^2+^ concentration is often detrimental for mitochondrial function (Rizzuto et al., 2012). Therefore, we assessed mitochondrial levels of Ca^2+^ and found that HKDC1-KO cells had significantly higher levels of Ca^2+^ (Fig 5F; left panel). We further confirmed this observation in a Hep3B2 cells where we used siRNAs to knockdown HKDC1 (Fig 5F; right panel). As higher Ca^2+^ levels in the mitochondria leads to decreased membrane potential (MMP) and induces reactive oxygen species (ROS) production we assessed both and we observed that HKDC1-KO cells had decreased MMP (as measured by tetramethyl rhodamine methyl ester (TMRM)) (Fig 5G) and enhanced ROS production (Fig 5H) (Bauer and Murphy, 2020, Giorgi et al., 2012, Ivanova et al., 2017, Calvo-Rodriguez et al., 2020). To investigate what could be the cause of increased mitochondrial Ca^2+^ in HKDC1-KO cells, we used transmission electron microscopy to investigate ER and mitochondrial morphology and their physical interaction upon HKDC1 ablation in HepG2 cells. We observed that HKDC1-KO cells displayed a markedly higher degree of ER apposition to mitochondria (Fig 5I). Hepatic mitochondrial content was also lower in HKDC1-KO cells than in EV cells, as measured by TEM (Fig 5J), and VDAC/COXIV expression (Supplementary Fig 5D). We also observed that mitochondria from HKDC1-KO cells were rounder and more swollen compared to the tubular mitochondria observed in EV cells (Fig 5I); however, a three-dimensional reconstruction of serial-sections TEMs would be a more definite way of confirming this observation. These data suggests that higher mitochondrial Ca^2+^ is associated with mitochondrial dysfunction in HKDC1-KO cells.

It has been shown by others that increased mitochondria-ER contact could be due to mitochondrial dysfunction, ER stress or both. We therefore, once again, analyzed our RNA-seq data for GO terms related to ER stress and observed that several (28 out of 30) significantly upregulated GO terms (biological process) in HKDC1-KO cells belonged to ER stress (Supplementary Fig 5E). We further assessed this finding using qPCR analysis for markers of ER stress and found that there was a significant upregulation of the ER chaperone protein BiP (binding-immunoglobulin protein aka GRP-78) in HKDC1-KO cells indicating elevated ER stress (Fig 5K) (Adams et al., 2019). We also found that in line with our RNA-seq data there was a significant upregulation of mRNA of activating transcription factor 4 (ATF4) and C/EBP-homologous protein (CHOP) which are the effectors of PKR-like ER kinase (PERK) (Fig 5K) (Adams et al., 2019). Furthermore, we also found elevated mRNA levels of inositol requiring enzyme 1α/β (IRE1) (Adams et al., 2019) (Fig 5K). IRE1 mediates its role in ER stress through multiple downstream proteins including XBP1 (X-box-binding protein 1) (Adams et al., 2019, Coelho and Domingos, 2014), but was not found to be altered in HKDC1-KO (Fig 5K). As PERK and IRE1 pathways are two of the three key unfolded protein response (UPR) signal activator protein pathways of ER stress (Adams et al., 2019), we explored the third arm of UPR, activating transcription factor 6 (ATF6); however, it was not found to be significantly altered upon HKDC1 ablation (Fig 5K) (Adams et al., 2019). UPR is elevated in response to ER stress to restore ER protein homeostasis as a primary mechanism for cell survival. However, if there is persistent activation of either of the three UPR pathways, caused by unmitigated severe ER stress, the cell decides to switch the signaling in favor of cell death (Adams et al., 2019, Coelho and Domingos, 2014, Wang et al., 2010, Hotamisligil and Davis, 2016). Therefore, we hypothesize that upon conditions where HKDC1 is not bound to the mitochondria (HKDC1-KO), there is increased mitochondria-ER interaction that leads to ER stress and mitochondrial dysfunction resulting in altered metabolism and cell death.

### Mitochondrial interaction of HKDC1 is essential for its role in HCC progression

Our studies have revealed how HKDC1 ablation in HCC produces multiple effects on HCC progression in part via inhibition of cell cycle progression and metabolic reprogramming where glucose consumption is enhanced but mitochondrial function is reduced. In combination, these effects result in reduction in cell proliferation and survival upon HKDC1 ablation. As shown by us (Pusec et al., 2019), HKDC1 contains a N-terminus targeting sequence similar to HK1 comprising the mitochondrial localization signal and we found that HKDC1 localizes at the mitochondria and interacts with mitochondrial protein such as VDAC. We therefore asked whether mitochondrial interaction of HKDC1 is necessary for its role in HCC metabolism, proliferation, and survival. To investigate this, we used an expression lentivirus that carried a truncated version of HKDC1 lacking the first 20 amino acids essential for mitochondrial binding (HKDC1-TR), as compared to our full length HKDC1 (HKDC1-FL). To avoid the involvement of endogenous HKDC1, we used our HKDC1-KO cells to overexpress HKDC1-TR and the full length HKDC1 (HKDC1-FL). Both lentiviruses produced robust mRNA and protein expression in HKDC1-KO cells (Supplementary Fig 6A-B). Moreover, we confirmed with a cell fractionation experiment that the HKDC1-TR did not localize at the mitochondria (Supplementary Fig 6C). Using these cells lines in proliferation, survival and metabolic assays, our data shows that HKDC1 ablation induced reduction in proliferation, survival and invasion which was significantly rescued by the HKDC1-FL overexpression but not by HKDC1-TR (Fig 6 A-C). Using Seahorse assay in these cell lines, HKDC1-FL overexpression was able to enhance both basal and maximal respiration compared to HKDC1-KO, whereas the HKDC1-TR overexpression remained similar to KO cells (Fig 6D). Moreover, in HepG2 cells, we observed a similar effect on mitochondrial complex activities where HKDC1-FL overexpression enhanced both complex I and III activities compared to KO cells and the TR version was similar to KO cells (Fig 6E). Finally, we checked for ER stress markers that were upregulated in HKDC1-KO cells and our data shows that the expression of these markers was restored to when HKDC1-FL cells but the expression either remained similar to HKDC1-KO cells or were further upregulated in HKDC1-TR cells (Fig 6F). These data confirm our hypothesis that mitochondrial interaction of HKDC1 is essential for its role in HCC.

**Fig 6.**
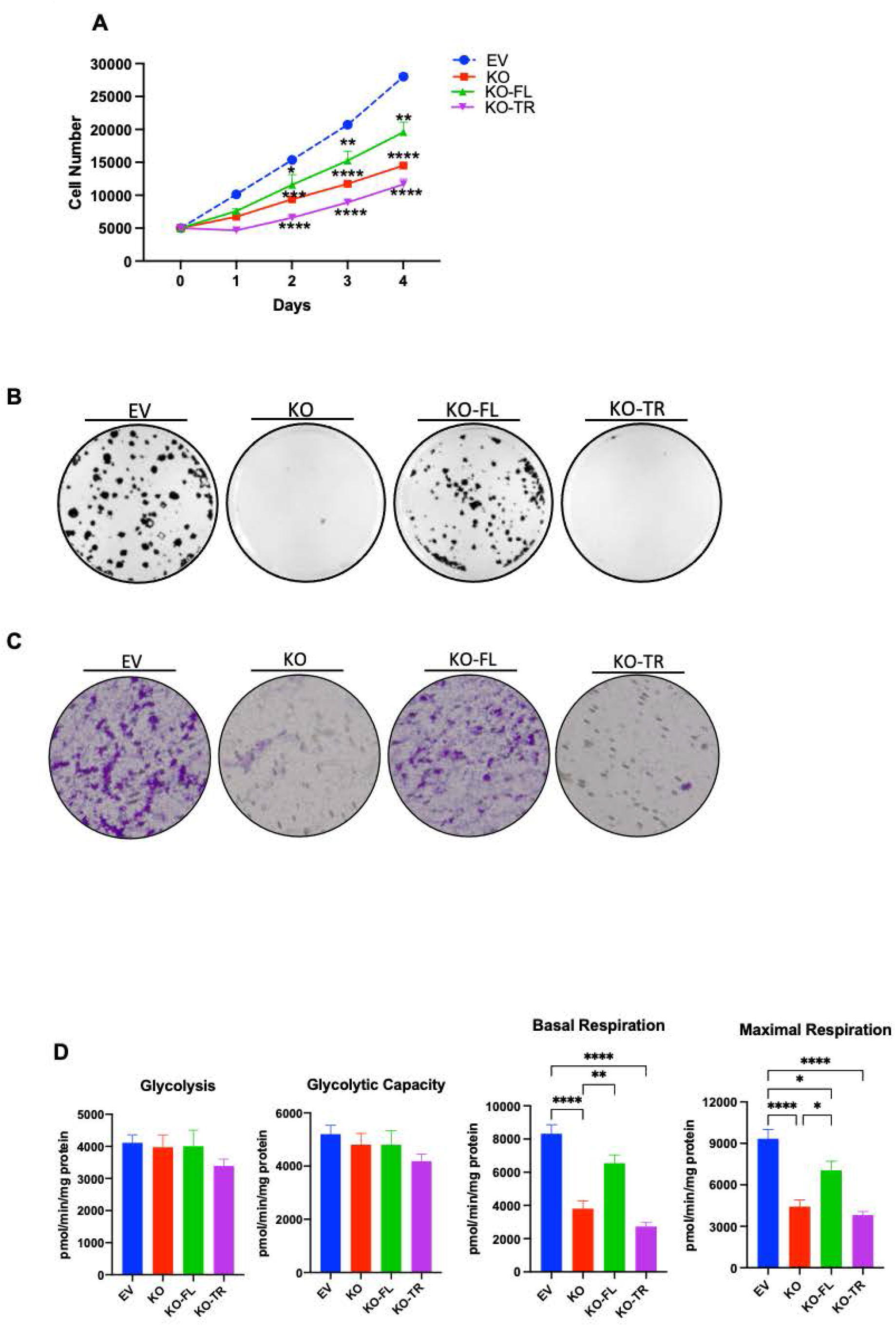
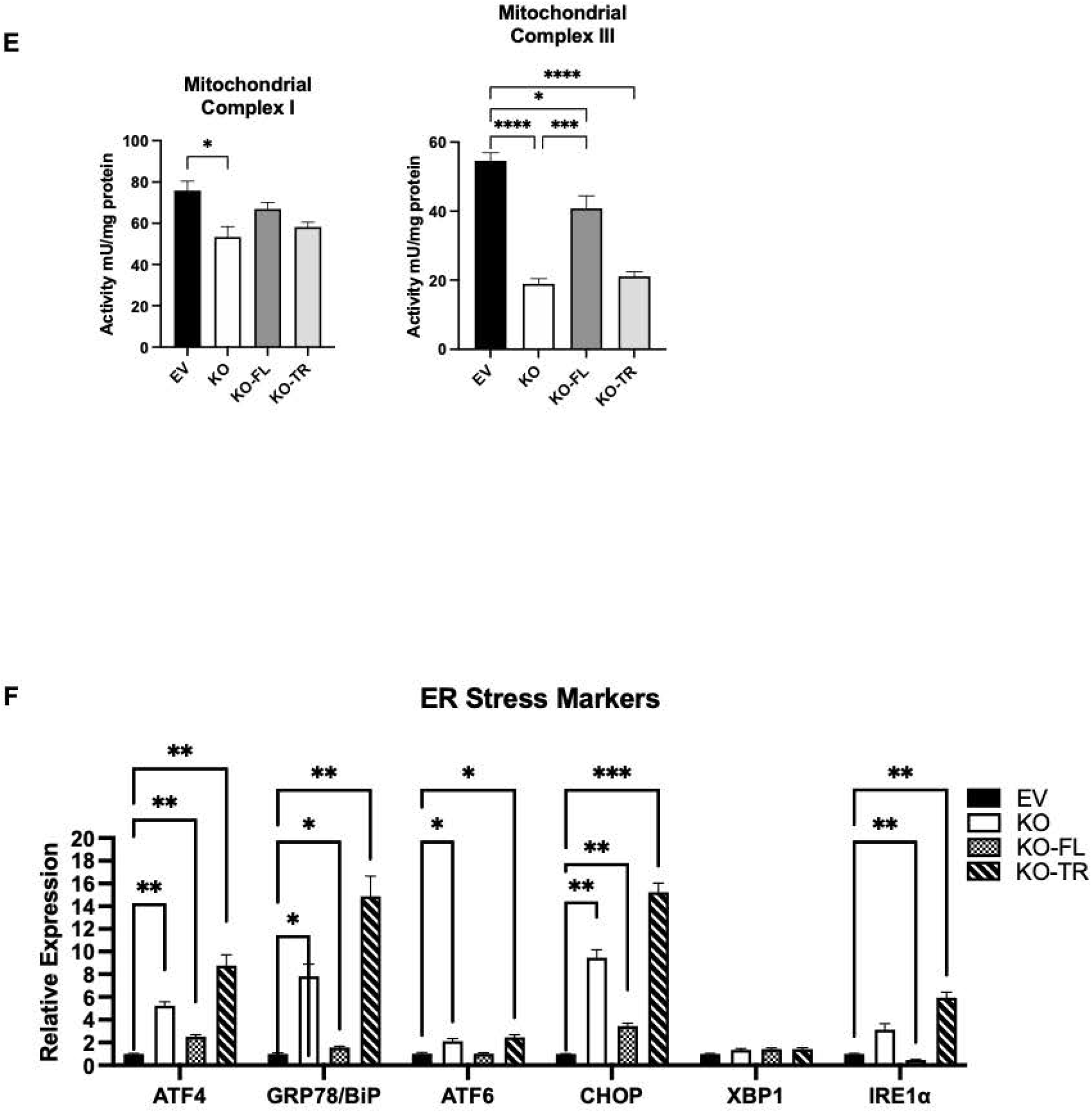
HKDC1-mitochondria interactions are necessary for mitochondrial function in HCC. Using the EV, HKDC1-KO, KO-FL and KO-TR cells, the following experiments were performed; **A)** cell proliferation assay, **B)** colony formation assay, **C)** transwell invasion assay, and **D)** seahorse metabolic analysis (ECAR and OCR). **E)** Activity assay for mitochondrial complex I (left panel) and complex II (right panel) for each cell lines was also performed. **F)** qPCR analysis to assess mRNA levels of ER stress markers. Values are ± SEM. All cell line experiments were performed 2-3 independent times with at least three replicates per experiments. Values are ± SEM; *p < 0.05; **p < 0.01; ***p < 0.001; ****p < 0.0001 by 2-way ANOVA.

## Discussion

Cancer cells undergo metabolic reprogramming by switching “on” or “off” the expression of specific genes to fulfill their catabolic and anabolic demands. As this is of extreme importance to the cancer cell, this distinction (from normal cells) can be potentially exploited to selectively target the cancer cells. HKs are the enzymes that catalyze the first step in glucose metabolism and are known to be altered in HCC (Smith, 2000). In the normal adult liver, this step is catalyzed by GCK (HK4), but expression of GCK is suppressed in HCC cells and instead we see a isoform switch to HK2 and HKDC1 (DeWaal et al., 2018, Zhang et al., 2016). HKDC1 expression is induced in a variety of human cancer types (Jiang et al., 2021, Smith, 2000) including HCC cells where it is minimally expressed in normal hepatocytes, thus, targeting it would selectively target cancer cells in HCC. Supporting this potential as a novel target, we found that HKDC1 deletion in human HCC cells inhibits proliferation and tumorigenesis both *in vitro* and *in vivo* and inhibits hepatocarcinogenesis in mice. We also show that HKDC1 interacts with the mitochondria in HCC cells and its deletion induces mitochondrial dysfunction. And finally, HKDC1’s role in HCC proliferation is due to its specific interaction with the mitochondria.

In this report, we show that HKDC1 ablation does not interfere with the expression of other HKs and most importantly, HK2 which is highly expressed in HCC. We and others have shown previously (Pusec et al., 2019, Chen et al., 2020) and here that HKDC1 has minimal hexokinase activity. As its ablation does not change cellular HK activity; we surprisingly observed a profound change increase in glucose uptake. With HKDC1 ablation, untargeted and targeted metabolomic experiments show increased glucose uptake with enhanced flux through the glycolysis shunt pathways PPP and HBP, with diminished TCA cycle metabolites. Mitochondrial binding of hexokinases has been suggested in cancer cells to be necessary for access of ATP derived from oxidative phosphorylation to efficiently phosphorylate glucose while others have shown this interaction to be antiapoptotic in nature (Robey and Hay, 2006, Cesar Mde and Wilson, 1998). We provide evidence for the first time that HKDC1 interaction with the mitochondria is essential to HCC, and its deletion induces mitochondrial dysfunction in HCC. HK2, which is the other highly expressed HK, has both cytosolic and mitochondrial localization. Moreover, HK2 is the major contributor to cellular HK activity and previously it has been shown that silencing HK2 leads to a decrease in glycolysis but does not induce mitochondrial dysfunction (DeWaal et al., 2018).

We and others have previously shown that HKDC1 binds to VDAC and regulates the permeability transition pore which provides a reasonable mechanism for HKDC1 mediated metabolic effects in HCC (Chen et al., 2020, Pusec et al., 2019). Our results provide evidence that HKDC1 ablation or silencing leads to decrease in mitochondrial function in HCC cells. We further show this by using a mitochondrial binding deficient HKDC1 (HKDC1-TR) which when re-expressed in HKDC1-KO cells cannot restore mitochondrial function. As cells make most of their energy in the form of ATP from mitochondrial function, HKDC1-KO cells had significantly less ATP than control cells. This decrease in cellular ATP might result in a metabolic stress where cells then resort to taking in more glucose to meet the energetic demands of the cell. However, since HKDC1-KO cells have impaired mitochondrial function, enhanced glucose flux resulting from increased uptake does not fuel the TCA cycle but is moved to glucose shunt pathways. As cancer cells in the synthetic (S) phase of the cell cycle need ATP to prepare the cells for division, a lack of ATP might result in cell cycle arrest as observed in HKDC1-KO cells. Therefore, in HKDC1 ablated cells, the resulting energetic crisis due to mitochondrial dysfunction results in inhibited proliferation of HCC cells.

ER stress is elevated in many cancers including HCC where it has been shown to play a pro-oncogenic role (Nakagawa et al., 2014, Wang et al., 2010, Yi et al., 2012) and inducing ER stress represents a new therapeutic strategy for killing tumor cells (Wang and Kaufman, 2014). HKDC1 has been previously shown to be a part of the cellular ER stress response (Evstafieva et al., 2018). It has been previously shown that mitochondrial dysfunction can induce ER stress due to reduced ATP levels in the cell (Liu et al., 2013). In this study, we found that HKDC1 ablation in HepG2 cells leads to more ER stress. If homeostasis is not restored, elevated ER stress triggers apoptosis or it could induce cell cycle arrest in the synthetic phase (Krebs et al., 2015, van Vliet and Agostinis, 2018, Szymański et al., 2017). Our data has shown that HKDC1-KO cells are arrested in the S-phase, and we hypothesize that HKDC1-KO mediated metabolic/energetic stress elevates ER stress which could be a probable mechanism explaining this cell-cycle blockade. Our data also shows that upon HKDC1 ablation, there is increased ER-mitochondria contact sites leading to mitochondrial Ca^2+^ overload and abnormal mitochondrial structure. Ca^2+^ regulates mitochondrial function in a complex manner which is incompletely understood where transient increase in mitochondrial Ca^2+^ can stimulate mitochondrial function (Rizzuto et al., 2012, Cárdenas et al., 2010). On the contrary, others have shown that increased ER-mitochondria contact results in increased mitochondrial Ca^2+^ accumulation which leads to mitochondrial dysfunction and increased ROS generation (Sebastián et al., 2012). Our data clearly shows that HKDC1-KO leads to mitochondrial dysfunction; however, future studies need to specifically address whether ER stress is a consequence of mitochondrial function or vice versa.

## Conclusion

Here, we have established HKDC1 as a novel contributor to the development and progression of HCC. Our studies could help explain why deletion of canonical HKs such as HK1 and 2 are not completely effective in inducing cell death where HKDC1 could be playing an additional role at the mitochondria. Specifically, HKDC1 deletion significantly affects glucose flux, energy metabolism and mitochondrial function. We also show that HKDC1 ablation results in less glucose flux to the TCA cycle leading to less cellular ATP generation and consequently, impacts cell cycle progression and ER stress induction. Our data shows that by interacting with mitochondria, HKDC1 regulates metabolism, proliferation, and survival of HCC. Moreover, because HKDC1 has nominal expression in normal hepatocytes, but is highly upregulated in HCC, novel small molecule and peptide-based inhibitors could be designed to specifically target HKDC1, in particular, its mitochondrial interaction in HCC.

## Acknowledgements

B.T.L. is supported by a Merit Review Award (I01BX00382), NIH grants (R01 DK104927, R01 DK111848, U01 DK127378, and P30 DK020595). MWK is supported by DOD Career Development Grant (W81XWH2010650). Liver cancer cell lines (SNU-387, 475) were kindly donated by Ron C. Gaba (Radiology & Pathology, UIC). Liver cancer cell lines (Hep3B2 and Huh7) were kindly donated by Dr. Nissim Hay (Department of Biochemistry and Molecular Genetics, UIC).

## Supplementary Fig Legends

**Supplementary Fig 1.**
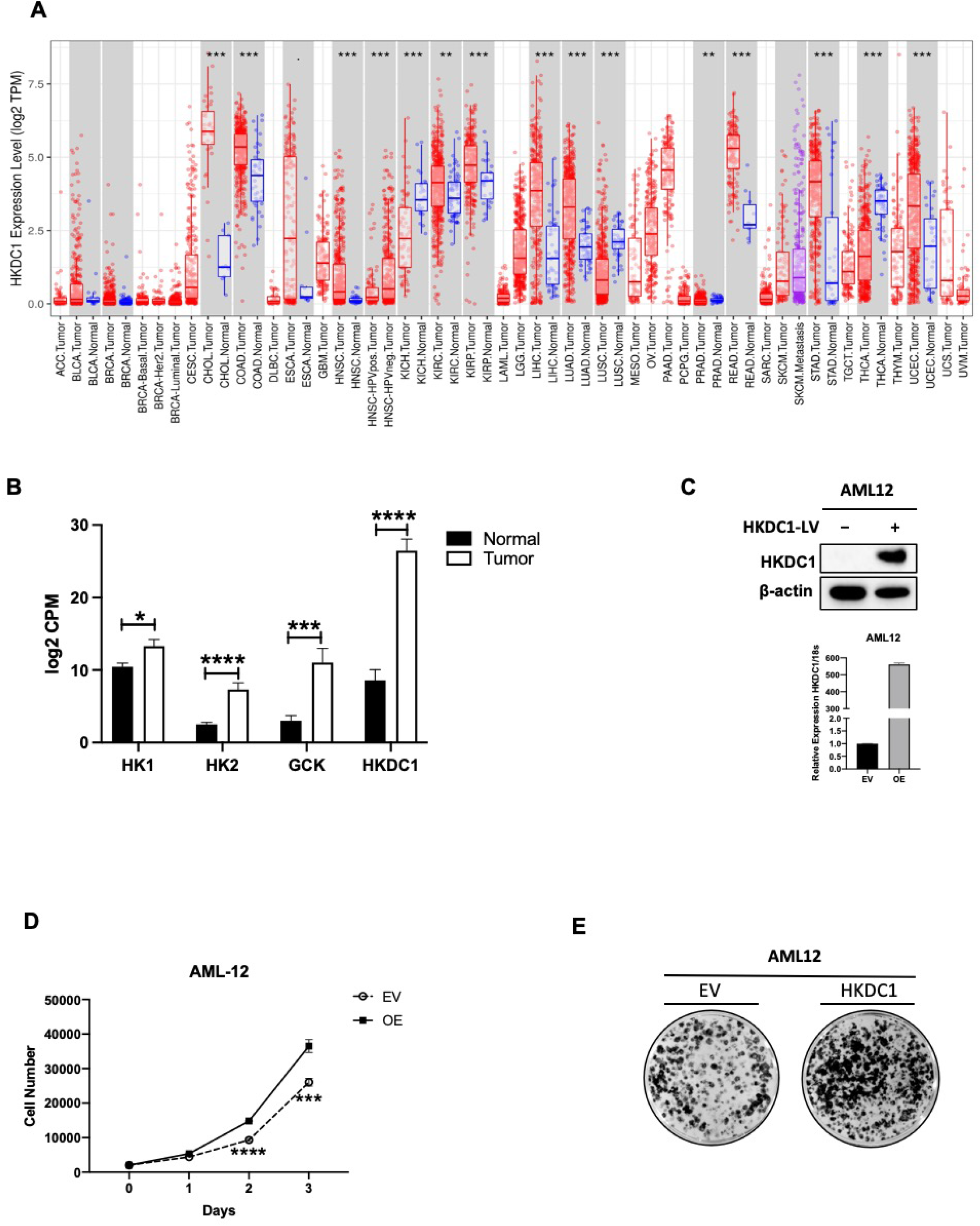
**A)** mRNA expression data of HKDC1 in various types of cancer in human patients were mined from the TCGA data set using the Timer website (https://cistrome.shinyapps.io/timer/). Supplementary Table 3 list all abbreviations for the tumor types included. **B)** Differential expression of HK1-2, GCK and HKDC1 HCC patients from TCGA data set (as described in methods) was mined; Tumor (n=369) and normal surrounding tissue (n=50). **C)** HKDC1 was overexpressed using lentivirus (LV) in AML12 cells; top panel (immunoblot) and bottom panel (qPCR) showing overexpressed HKDC1 protein and mRNA levels. **D)** cell proliferation curves in modulated AML12 cells, EV: cells with empty vector; OE: cells overexpressing HKDC1. **E)** colony forming assay, EV: cells with empty vector; HKDC1: cells overexpressing HKDC1. All cell line experiments (**C-E**) were performed 2-3 times with 3-8 replicates per experiment. Values are ± SEM; **p < 0.01; ***p < 0.001; ****p < 0.0001 by Wilcoxon signed-rank test (for 2A), 2-way ANOVA (for 2B) and students t-test (for 2D).

**Supplementary Fig 2.**
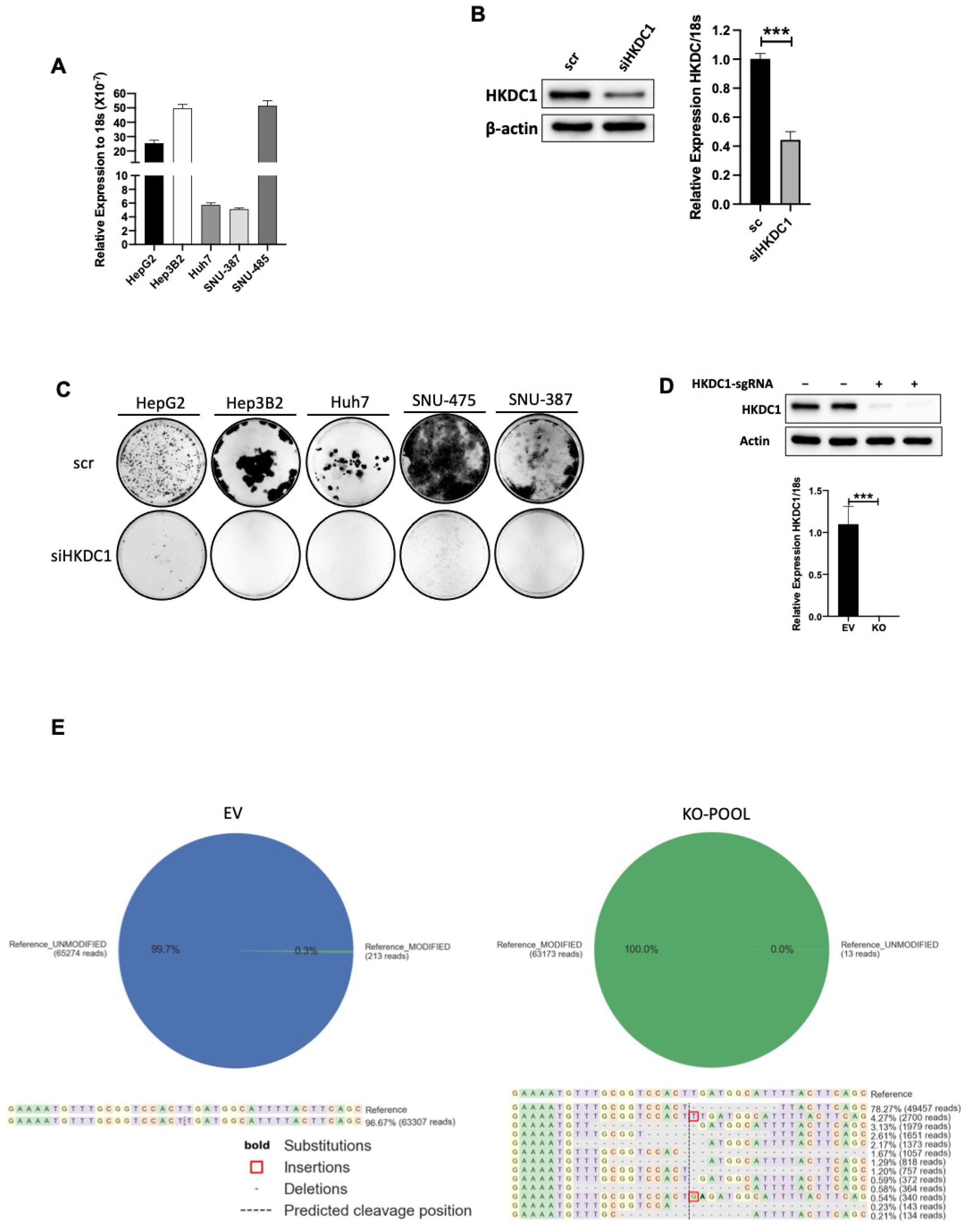
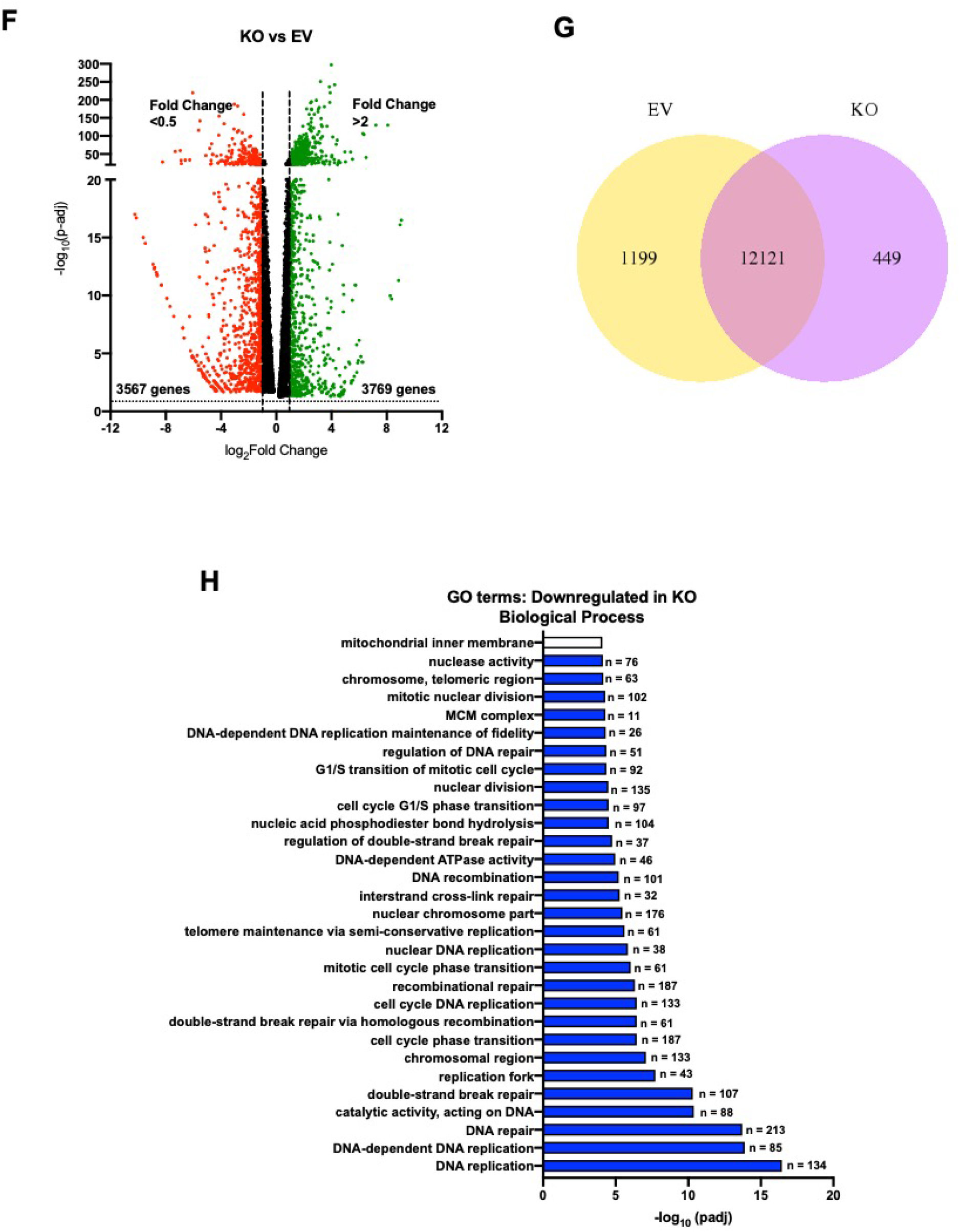
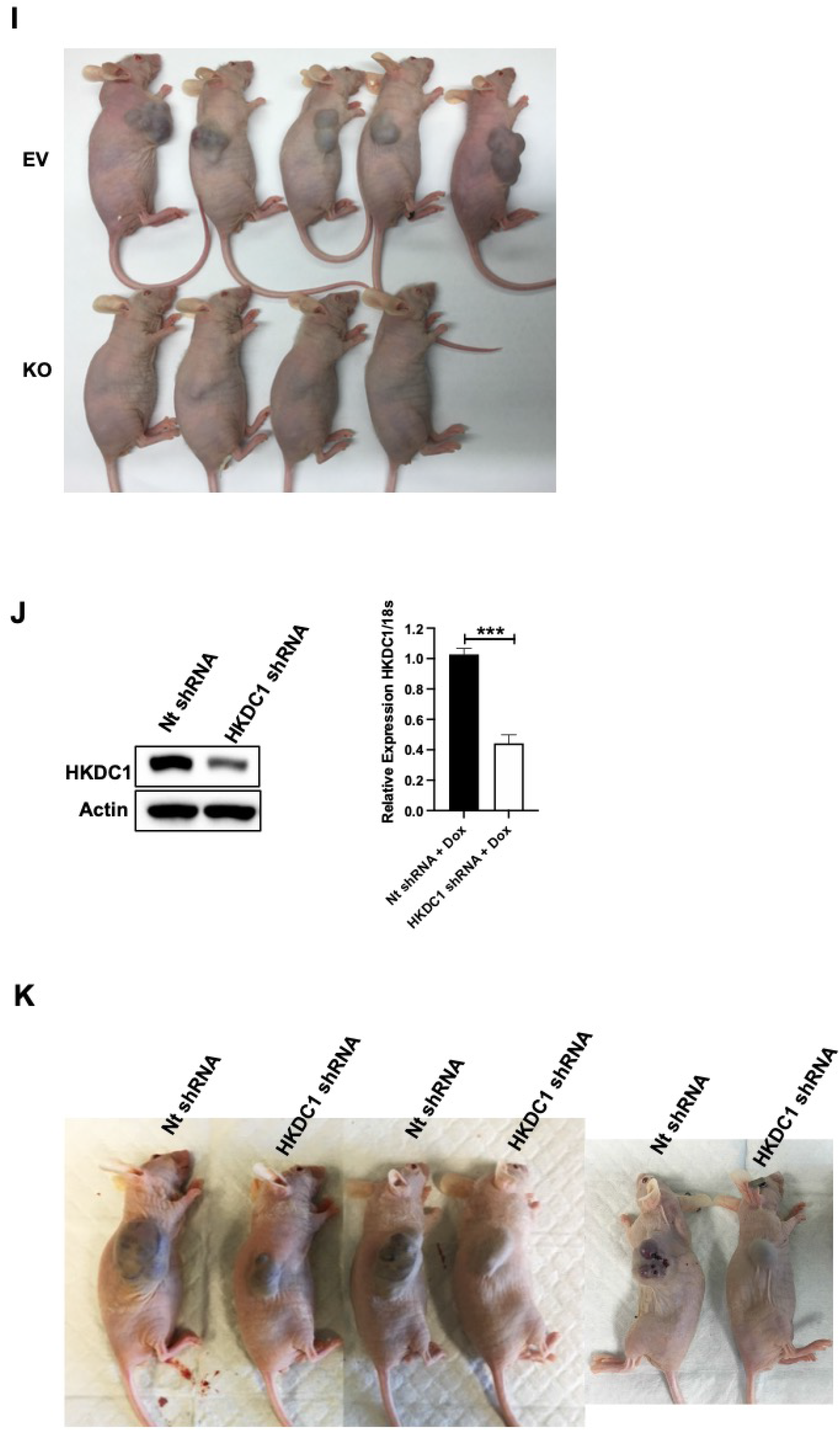
**A)** HKDC1 mRNA expression in a panel of human HCC cell lines as compared to expression of 18s in each cell line. **B)** siRNA mediated HKDC1 knockdown in HepG2 cells. Cell were transfected with siRNAs for 48h following which cells were harvested and subjected to immunoblot (left panel) and qPCR (right panel) to assess HKDC1 protein and mRNA levels. **C)** colony forming assay in the panel of HCC cell lines after siRNA mediated knockdown of HKDC1 **D)** Crispr-Cas9 was used to knockout HKDC1 in HepG2 cells; top panel shows protein expression of HKDC1 after sgRNA transfection and bottom panel shows mRNA expression. **E)** Genomic sequencing in the KO pool (see methods) was done to provide evidence for Crispr-cas9 mediated genomic alteration of *HKDC1* (left panel EV; right panel KO). **F)** Volcano plot created from list of differentially expressed genes from RNA-seq data. **G)** Unique and overlapping expressed genes in EV and KO cells (from RNA-seq data analysis). **H)** Gene ontology (GO) terms plotted for downregulated GO terms (from RNA-seq data analysis). **I)** *in vivo* tumor growth. 1X10^6^ EV or KO cells were inoculated into mice (n=4-5). Images of mice bearing tumors were taken at endpoint. **J)** Hep3B2 cells were transfected with shHKDC1 or (non-target) ntshRNA and transfected cells were selected with appropriate antibiotics. Cells were then treated with doxycycline for 3 days and HKDC1 protein and mRNA expression was assessed by immunoblot (left panel) and qPCR (right panel). **K)** Hep3B2 cells were transfected with shHKDC1 or (non-target) ntshRNA and transfected cells were selected with appropriate antibiotics, 1X10^6^ cells were inoculated into mice (n=4). When tumors were visible, mice were given doxycycline (in diet) for 7 days to activate shRNAs. Tumor growth was measured weekly till 8 weeks after appearance of tumor with a vernier caliper. Images of mice bearing tumors were taken at endpoint. All cell line experiments (**A-D**) were performed 2-3 times with 3-8 replicates per experiment. Values are ± SEM; ***p < 0.001; ****p < 0.0001 students t=test (for 2D) and 2-way ANOVA (for 2B, J).

**Supplementary Fig 3.**
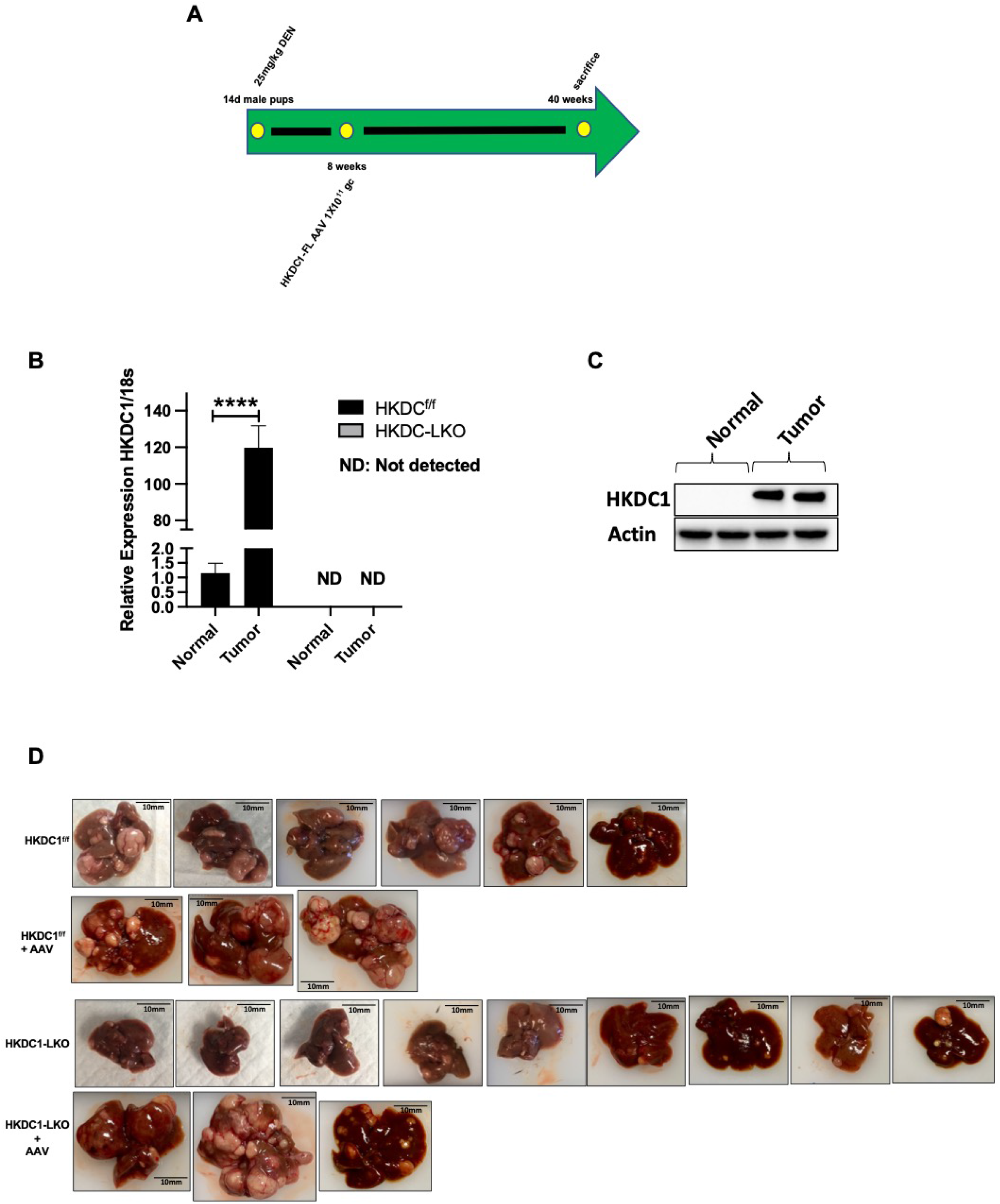
**A)** Representative images from DEN induced tumorigenesis experiment. Two-week-old HKDC1^f/f^ and HKDC1-LKO male mice were injected with DEN (25 mg/kg). When mice were 8 weeks old, both groups were further divided into groups where one group received AAV expressing human HKDC1 (HKDC1^f/f^+AAV and HKDC1-LKO+AAV) and the AAV expressing null vector was used as the control with the two other groups (HKDC1^f/f^ and HKDC1-LKO), with N=4-7 per group. **B)** Scheme for DEN treated mice. 14d male pups were injected with 25m/kg DEN intra-peritoneally. When mice were 8 weeks old, mice form both groups were further divided into groups where one group received AAV-Null and the other AAV-HKDC1-FL. Mice were monitored till 40 weeks and sacrificed at that time point. **C)** mRNA expression of *HKDC1* from normal and tumor from liver tissue (n=3). **D)** immunoblot to assess protein expression of HKDC1 in normal and tumor samples of HKDC1^f/f^ mice liver sample (n=2). Values are ± SEM; ****p < 0.0001 by 2-way ANOVA.

**Supplementary Fig 4.**
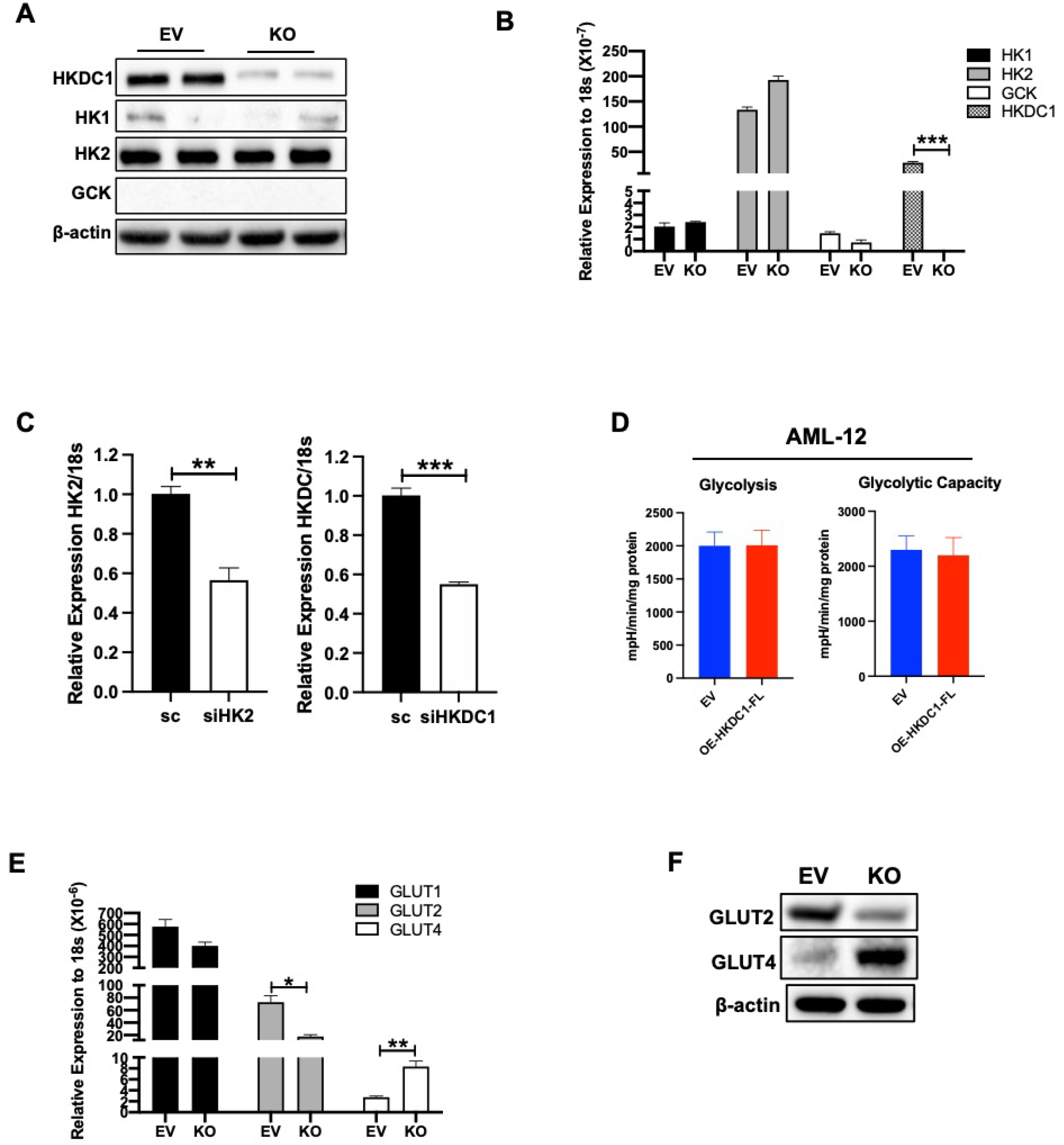
EV and KO cells were analyzed for expression of different HKs by **A)** immunoblot and **B)** qPCR. **C)** HepG2 cells were treated with siRNA against either HK2 or HKDC1 for 24h, cells were lysed, qPCR analysis (relative expression to scramble siRNA (scr)) showing siRNA efficiency. **D)** Seahorse metabolic analysis (ECAR) of AML cells expressing either empty vector (EV) or full-length HKDC1 (OE-HKDC1-FL). **E)** qPCR analysis in EV and KO cells showing the expression level of *GLUT1*, *GLUT2* and *GLUT4*. **F)** immunoblot showing expression of GLUT2 and GLUT4 in EV and KO cells. All cell line experiments were performed 2-3 times with 3-5 replicates per experiment. Values are ± SEM; **p < 0.01; ***p < 0.001 by student’s t-test (for 4C) or 2-way ANOVA (for 4B and E).

**Supplementary Fig 5.**
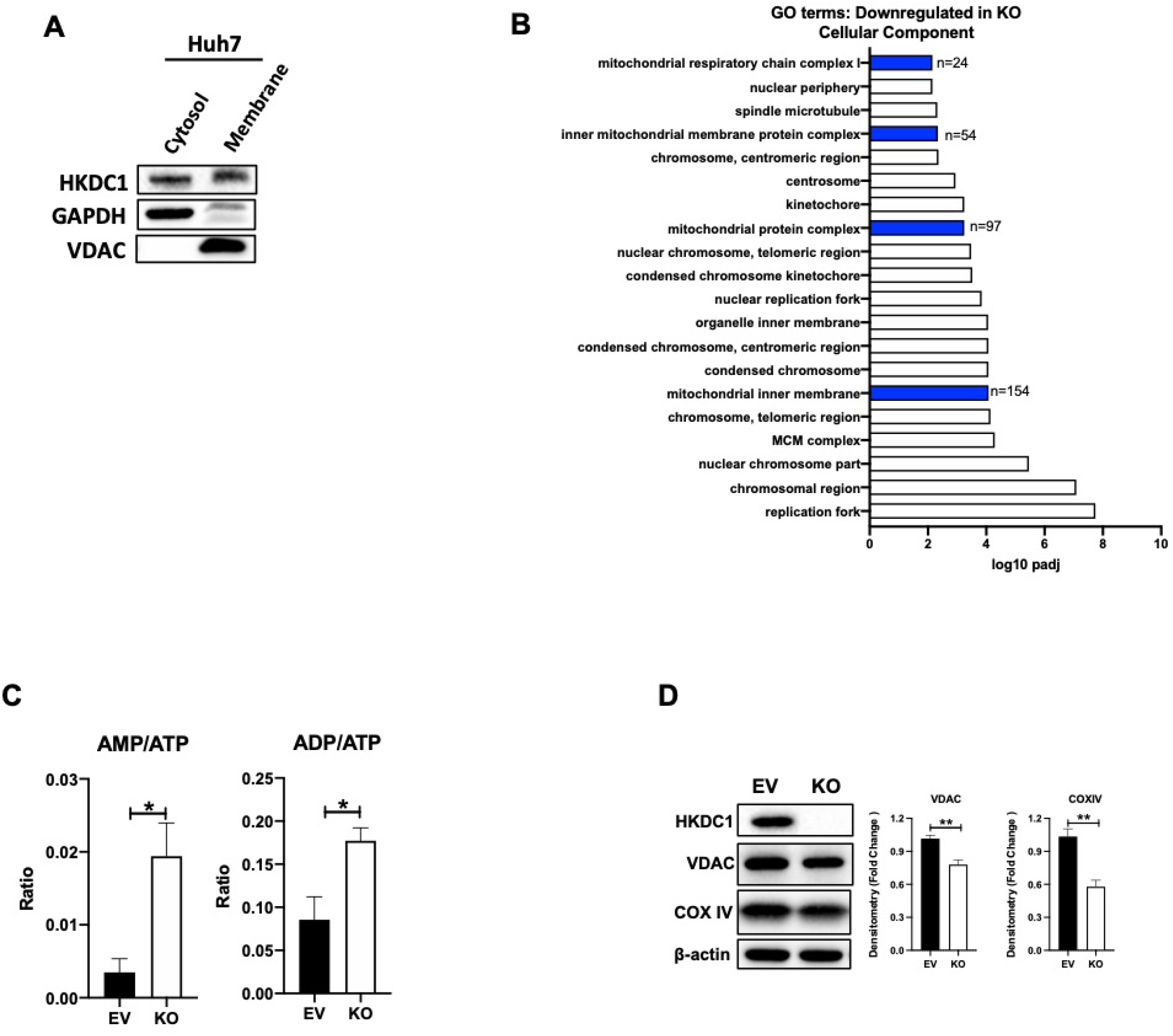
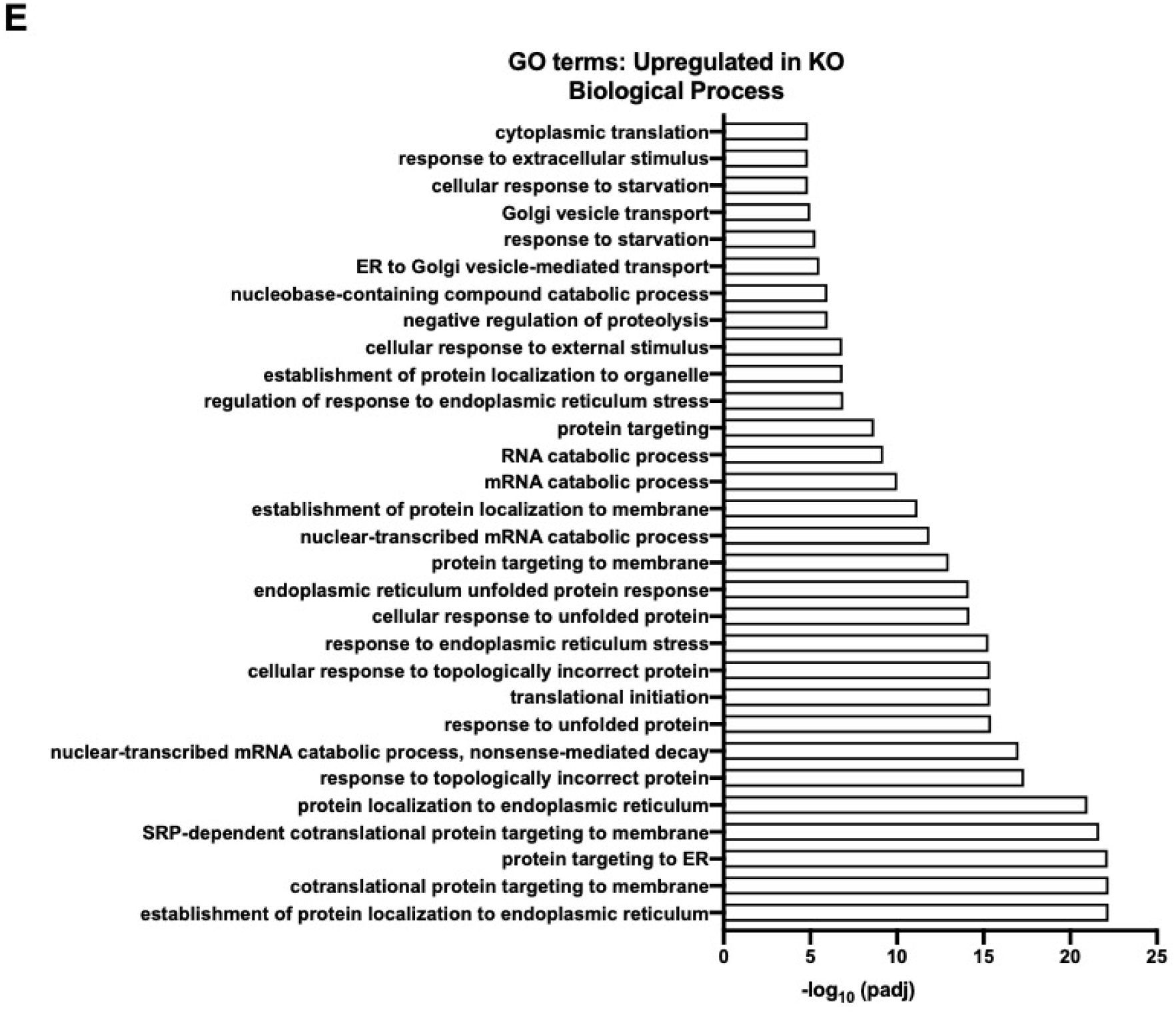
**A)** Subcellular fractionation of HuH7 cells lines followed by immunoblot to show HKDC1 expression in cytosolic and membrane compartments. **B)** Gene ontology (GO) terms plotted for significantly downregulated GO terms in cellular component (from RNA-seq data analysis). **C)** Ratio of AMP/ATP and ADP/ATP values from steady-state metabolomics (n=3). **D)** immunoblot of EV and KO showing expression of mitochondrial proteins; right panels show densitometry (immunoblot is representative of 3 different blots). **E)** Gene ontology (GO) terms plotted for significantly upregulated GO terms in biological processes (from RNA-seq data analysis). Values are ± SEM; *p < 0.05; **p < 0.01 by student’s t-test.

**Supplementary Fig 6.**
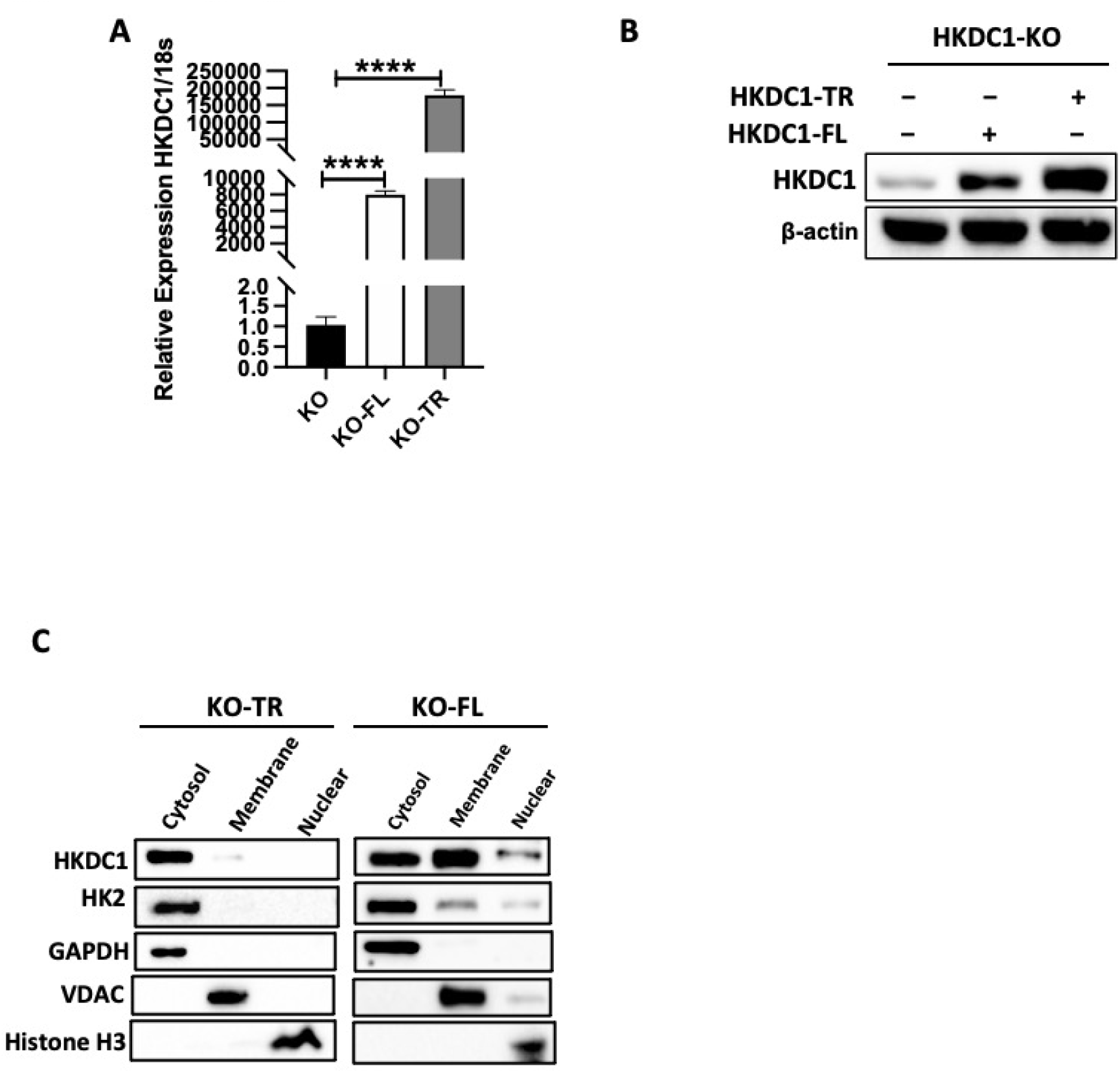
HKDC1-KO (KO) HepG2 cells were used to overexpress either HKDC1-FL (KO-FL) or HKDC1-TR (KO-TR) by lentiviruses **A)** mRNA expression and **B)** protein expression was analyzed after cells stably transduced cells were selected with appropriate antibiotics. **C)** Subcellular fractionation of KO-FL and KO-TR cells followed by immunoblot to show HKDC1 expression in cytosolic and membrane compartments. Values are ± SEM; **p < 0.01; ***p < 0.001, ****p < 0.0001 by one-way ANOVA (for 6A, D)

**Supplementary Table 1:**
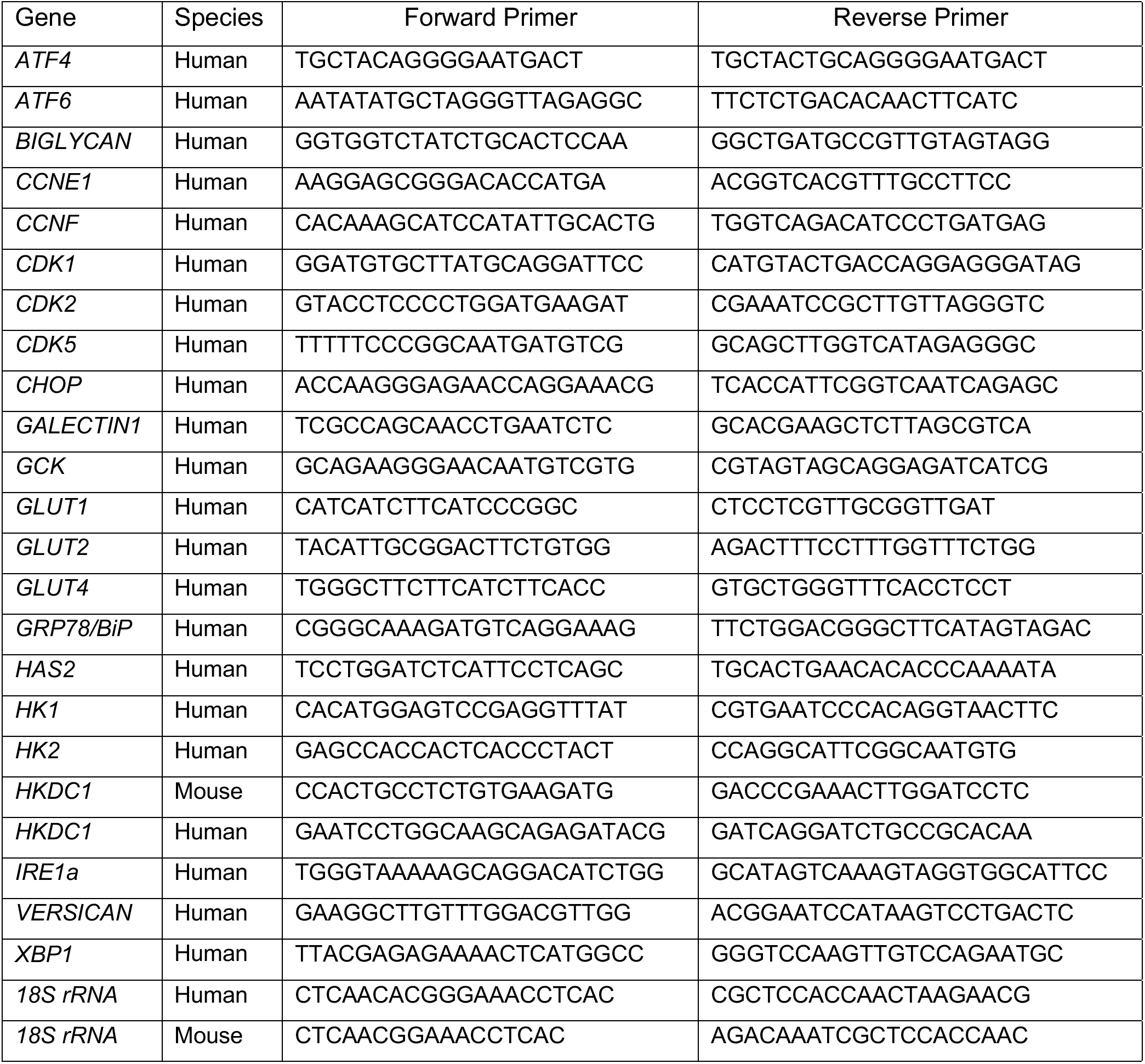
List of Primers.

**Supplementary Table 2:** List of Antibodies.

| Antibody | Manufacturer | Cat No | Host | Dilution* |
| --- | --- | --- | --- | --- |
| Beta-actin | Sigma | A2228 | Mouse | 1:5000 |
| COXIV | Cell Signaling Technology | 4850 | Rabbit | 1:1000 |
| GAPDH | Cell Signaling Technology | 14C10 | Rabbit | 1:2000 |
| GCK | Santa Cruz Biotechnology | sc-17819 | Mouse | 1:1000 |
| Glut 2 | Novus Biologicals | NBP2-22218 | Rabbit | 1:1000 |
| Glut 4 | Cell Signaling Technology | 2213 | Mouse | 1:1000 |
| Histone H3 | Cell Signaling Technology | 4499 | Rabbit | 1:1000 |
| HK1 | Cell Signaling Technology | 2024 | Rabbit | 1:1000 |
| HK2 | Cell Signaling Technology | 2867 | Rabbit | 1:1000 |
| HKDC1 | Abcam | ab228729 | Rabbit | 1:1000 |
| VDAC | Cell Signaling Technology | 4661 | Rabbit | 1:1000 |
| Anti-Rabbit | Cell Signaling Technology | 7074 | Goat | 1:10000 |
| Anti-Mouse | Cell Signaling Technology | 7076 | Horse | 1:10000 |
| *All dilutions were made in phosphate buffered saline with 5% BSA and 0.1% Tween 20 |  |  |  |  |

**Supplementary Table 3:** TCGA study abbreviations.

|  |  |
| --- | --- |
| ACC | Adrenocortical carcinoma |
| BLCA | Bladder Urothelial Carcinoma |
| LGG | Brain Lower Grade Glioma |
| BRCA | Breast invasive carcinoma |
| CESC | Cervical squamous cell carcinoma and endocervical adenocarcinoma |
| CHOL | Cholangiocarcinoma |
| LCML | Chronic Myelogenous Leukemia |
| COAD | Colon adenocarcinoma |
| CNTL | Controls |
| ESCA | Esophageal carcinoma |
| FPPP | FFPE Pilot Phase II |
| GBM | Glioblastoma multiforme |
| HNSC | Head and Neck squamous cell carcinoma |
| KICH | Kidney Chromophobe |
| KIRC | Kidney renal clear cell carcinoma |
| KIRP | Kidney renal papillary cell carcinoma |
| LIHC | Liver hepatocellular carcinoma |
| LUAD | Lung adenocarcinoma |
| LUSC | Lung squamous cell carcinoma |
| DLBC | Lymphoid Neoplasm Diffuse Large B-cell Lymphoma |
| MESO | Mesothelioma |
| MISC | Miscellaneous |
| OV | Ovarian serous cystadenocarcinoma |
| PAAD | Pancreatic adenocarcinoma |
| PCPG | Pheochromocytoma and Paraganglioma |
| PRAD | Prostate adenocarcinoma |
| READ | Rectum adenocarcinoma |
| SARC | Sarcoma |
| SKCM | Skin Cutaneous Melanoma |
| STAD | Stomach adenocarcinoma |
| TGCT | Testicular Germ Cell Tumors |
| THYM | Thymoma |
| THCA | Thyroid carcinoma |
| UCS | Uterine Carcinosarcoma |
| UCEC | Uterine Corpus Endometrial Carcinoma |
| UVM | Uveal Melanoma |

